# Recruitment of Polo-like kinase couples synapsis to meiotic progression via inactivation of CHK-2

**DOI:** 10.1101/2021.06.20.449183

**Authors:** Liangyu Zhang, Weston T. Stauffer, John S. Wang, Fan Wu, Zhouliang Yu, Chenshu Liu, Abby F. Dernburg

## Abstract

Meiotic chromosome segregation relies on synapsis and crossover recombination between homologous chromosomes. These processes require multiple steps that are coordinated by the meiotic cell cycle and monitored by surveillance mechanisms. In diverse species, failures in chromosome synapsis can trigger a cell cycle delay and/or lead to apoptosis. How this key step in “homolog engagement” is sensed and transduced by meiotic cells is unknown. Here we report that in *C. elegans*, recruitment of the Polo-like kinase PLK-2 to the synaptonemal complex triggers phosphorylation and inactivation of CHK-2, an early meiotic kinase required for pairing, synapsis, and double-strand break induction. Inactivation of CHK-2 ends double-strand break formation and promotes crossover designation and cell cycle progression. These findings illuminate how meiotic cells ensure crossover formation and accurate chromosome segregation.

**Summary:** Accurate chromosome segregation during meiosis requires crossovers between each pair of homologs. Zhang *et al*. show that meiotic progression in *C. elegans* involves inactivation of CHK-2 by PLK-2 in response to synapsis and formation of crossover precursors on all chromosomes.

## Introduction

Meiotic progression differs in many ways from a mitotic cell cycle. A single round of DNA replication at meiotic entry is followed by two nuclear divisions. Between replication and the first division is an extended period known as meiotic prophase, during which chromosomes pair, synapse, and recombine to establish physical links (chiasmata) between each pair of homologs. Together with sister chromatid cohesion, chiasmata direct bipolar orientation and segregation of homologous chromosomes during Meiosis I (Hillers et al., 2017; Hunter, 2015; Ur and Corbett, 2021).

During early meiosis, double-strand breaks (DSBs) are induced and resected. These intermediates undergo strand invasion to establish joint molecules between homologs, which recruit a set of factors collectively known as “pro-crossover” or “ZMM” proteins. Many of these intermediates eventually lose the ZMM proteins and are resolved as non-crossovers, while the subset that retains these factors is “designated” to be resolved as crossovers (COs) (Hunter, 2015; Pyatnitskaya et al., 2019).

In *C. elegans*, formation and initial processing of DSBs requires the activity of CHK-2, a meiosis-specific homolog of the DNA damage transducing kinase Chk2/CHEK2 (MacQueen and Villeneuve, 2001; Oishi et al., 2001). CHK-2 is eventually inactivated at mid-pachytene; this coincides with the end of DSB formation and the resolution of many intermediates through a “generic” (as opposed to meiosis-specific) homologous recombination pathway. A similar transition during mid-prophase has been described in budding yeast and mammals. In each case this progression depends on assembly of the synaptonemal complex (SC), a meiosis-specific protein scaffold that assembles between homologous chromosomes (Enguita-Marruedo et al., 2019; Hayashi et al., 2007; Kauppi et al., 2013; Lee et al., 2021; Mu et al., 2020; Murakami et al., 2020; Nadarajan et al., 2017; Thacker et al., 2014; Wojtasz et al., 2009).

CHK-2 becomes active upon meiotic entry. The mechanism of activation has not been fully established, but it is promoted by targeted degradation of a CHK-2 inhibitor, PPM-1.D/Wip1 (Baudrimont et al., 2022). Work in budding yeast has shown that a CHK-2 ortholog, Mek1, undergoes trans-autophosphorylation of its activation loop (Carballo et al., 2008; Niu et al., 2007); this mechanism is likely conserved in *C. elegans* since the activation loop of CHK-2 also contains a CHK-2 consensus phosphorylation motif (R-x-x-S/T) (O’Neill et al., 2002). Like all Chk2 homologs, CHK-2 contains a forkhead-associated (FHA) domain and thus recognizes phosphorylated motifs of the consensus (pT-X-X-[I/L/V]) (Durocher and Jackson, 2002; Li et al., 2000). While mammalian Chk2 undergoes homodimerization through binding of its FHA domain to N-terminal motifs phosphorylated by ATM (Oliver et al., 2006), the meiosis-specific Mek1 and CHK-2 kinases lack this regulatory domain. Trans-activation is promoted by binding at high density to target motifs on other proteins, such as Hop1 in budding yeast (Carballo et al., 2008). In particular, the Pairing Center proteins HIM-8 and ZIM-1, -2, -3 contain such motifs and recruit CHK-2 during early meiotic prophase. By phosphorylating a Polo-box interacting motif on these proteins, CHK-2 primes the recruitment of the Polo-like kinase PLK-2 (Harper et al., 2011; Kim et al., 2015). Together, CHK-2 and PLK-2 at Pairing Centers modify NE proteins including lamin (LMN-1) and the LINC protein SUN-1, which promotes chromosome pairing and synapsis (Link et al., 2018; Penkner et al., 2009; Sato et al., 2009; Woglar et al., 2013). Thus, CHK-2 is required for chromosome pairing as well as for DSB induction.

By mid-pachytene, most of the early recombination intermediates disappear, and six “designated crossover” foci, which show strong localization of the cyclin homolog COSA-1 and other pro-CO factors, are detected in each nucleus, one per chromosome pair (Woglar and Villeneuve, 2018; Yokoo et al., 2012; Zhang et al., 2018). This accumulation of pro-CO proteins at the sites that will eventually become crossovers is termed “crossover designation.” Collectively, pro-CO factors are thought to prevent non-crossover resolution and eventually promote the resolution of these late intermediates as crossovers. The mechanism and timing of CO resolution in *C. elegans* remain somewhat unclear.

Defects in synapsis, or absence of any of the factors required to establish crossover intermediates, results in activation of a “crossover assurance checkpoint” and a delay in cell cycle progression (reviewed in Yu et al., 2016). Activation of the checkpoint is detected cytologically as an extended region of CHK-2 activity and a delay in the appearance of designated crossover sites (Kim et al., 2015; Rosu et al., 2013; Stamper et al., 2013; Woglar and Villeneuve, 2018; Zhang et al., 2018). Defects in synapsis result in higher sustained levels of CHK-2 than defects in recombination (Kim et al., 2015). This feedback regulation requires a family of HORMA domain proteins that bind to the chromosome axis (Kim et al., 2015), but how cells detect defects in synapsis or crossover intermediates and how CHK-2 activity is prolonged remain unknown.

## Results

### CHK-2 inhibits CO designation

To monitor the activity of CHK-2 in *C. elegans* oocytes (Figure 1A), we used a phospho-specific antibody that recognizes CHK-2–dependent phosphorylation sites on HIM-8 and ZIM-1/2/3 at Pairing Centers (Kim et al., 2015). Co-staining of pHIM-8/ZIMs and GFP::COSA-1 confirmed that crossover designation occurs concomitantly with loss of CHK-2 activity (Figure 1B-D). Both CHK-2 inactivation and the CO designation are delayed in mutants that activate the crossover assurance checkpoint (Kim et al., 2015; Zhang et al., 2018). We thus wondered whether CHK-2 activity inhibits CO designation. Because CHK-2 activity during early meiotic prophase is essential for DSB formation and synapsis (MacQueen and Villeneuve, 2001; Oishi et al., 2001), and thus for the establishment of recombination intermediates that are prerequisite for CO designation, we exploited the auxin-inducible degradation (AID) system to deplete CHK-2 activity (Nishimura et al., 2009; Zhang et al., 2015). The activity of degron-tagged CHK-2 became undetectable in meiotic nuclei within 3 hours of auxin treatment (Figure 2– figure supplement 1A-B). Depletion of CHK-2 shifted the appearance of nuclei containing bright COSA-1 foci to a more distal (earlier) position in the germline, indicative of accelerated CO designation and cell cycle progression (Figure 2A and C).

**Figure 1.**
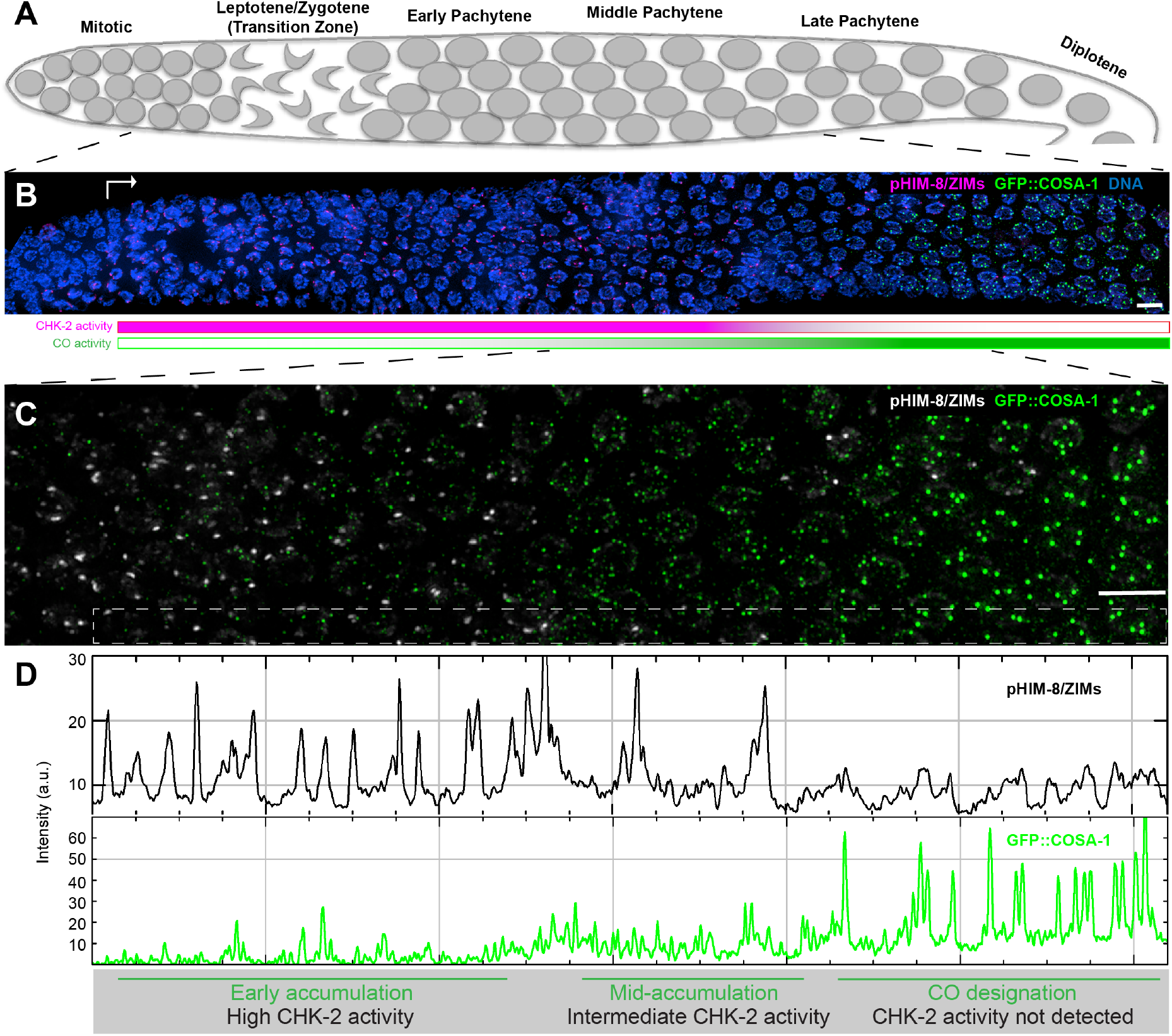
Temporal profile of CHK-2 activity and crossover designation during meiotic prophase in *C. elegans*. (A) Schematic of meiotic prophase I in the *C. elegans* hermaphrodite germline. Meiotic progression is readily visualized in *C. elegans* due to the simple organization of the germline. Cells enter meiosis in the distal “arms” of the gonad and move proximally towards the spermatheca and uterus with a velocity of about one row per hour (Deshong et al., 2014). Homolog pairing and synapsis and DSB induction occur during the first few hours following meiotic entry, in the “transition zone” region corresponding to leptotene and zygotene. DSBs were then processed to form recombination intermediates in early pachytene. In mid-pachytene, only one recombination intermediate on each pair of chromosomes is “designated” as an eventual crossover site. (B) Images of representative prophase nuclei from meiotic onset to late pachytene show the crossover intermediates (indicated by GFP::COSA-1, green) and CHK-2 activity (indicated by phospho-HIM-8 and -ZIM-1/2/3 immunofluorescence, magenta). Bright GFP::COSA-1 foci appear following inactivation of CHK-2. White arrow indicates meiotic onset. Scale bars, 5 µm. (C) Enlargement of the mid-pachytene nuclei shown in (B). phospho-HIM/ZIMs was pseudo colored to white for easy observation. Scale bars, 5 µm. (D) Plot profile of phospho-HIM-8/ZIMs immunofluorescence (upper) and GFP::COSA-1 (lower) intensity in the boxed region indicated in (C). Accumulation of pro-CO proteins can be stratified into three stages: an early stage with high CHK-2 activity and variable numbers of dim GFP::COSA-1; an intermediate stage with brighter and more visible GFP::COSA-1 foci, and a post-designation stage, with a single CO-designated sites per chromosome pair marked by bright GFP::COSA-1 fluorescence. The intensity of bright GFP::COSA-1 foci is relatively constant upon formation until the end of pachytene. According to this property, we determined the bright COSA-1 foci. In this work, we used the appearance of bright COSA-1 foci (corresponding to CO designation) to evaluate the timing of crossover activity accumulation and the corresponding cell cycle transition.

**Figure 2.**
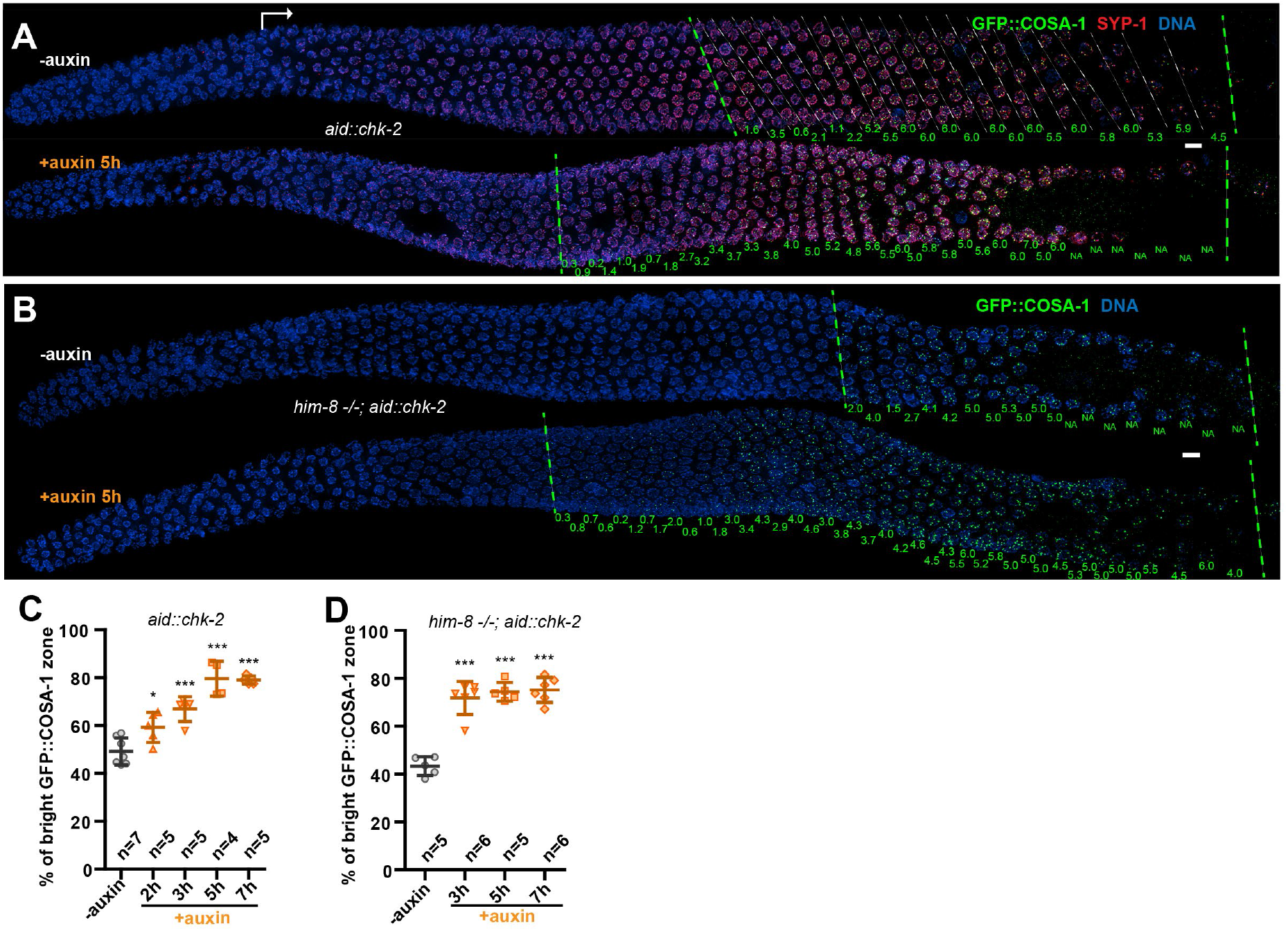
CHK-2 inhibits CO designation. (A) Representative hermaphrodite gonads stained for GFP::COSA-1 (green), SYP-1 (red) and DNA (blue). Nuclei with bright COSA-1 foci are observed at a more distal position following CHK-2 depletion. Worms were exposed to 1 mM auxin (or 0.25% ethanol lacking auxin) for 5 h before fixation. Dashed green lines indicate the earliest nuclei with bright COSA-1 foci and the end of pachytene, respectively. White arrow indicates meiotic onset. The average number of bright COSA-1 foci per nucleus in each row is indicated below each image in green. Scale bars, 5 µm. (B) Germline from a *him-8* mutant hermaphrodite stained for GFP::COSA-1 (green), and DNA (blue), showing early appearance of bright COSA-1 foci upon CHK-2 depletion. Worms were treated with 1 mM auxin or solvent (0.25% ethanol) control for 5 h before analysis. Scale bars, 5 µm. Note: the same image with RAD-51 is shown in Figure 2– figure supplement 1C. (C) CO designation occurs earlier upon CHK-2 depletion, as described in (A). Worms were exposed to auxin (or solvent control) for 2, 3, 5 or 7 h before analysis. We define the “Bright GFP::COSA-1 zone” as the length of the region from where bright GFP::COSA-1 foci appear to the end of pachytene, before oocytes form a single row of cells. Since the length of each meiotic stage region varies among individual animals, while the ratio between stages is relatively constant. We used the ratio of the length of “Bright GFP::COSA-1 zone” to the length of the region from meiotic onset to the end of pachytene to reflect the timing of bright GFP::COSA-1 foci appearance and crossover designation. To simplify, we hereafter use “% of bright GFP::COSA-1 zone” in graphs to indicate this ratio. Meiotic onset was determined by the staining of meiosis-specific proteins SYP-1 and/or HTP-3. *n* = number of gonads scored for each condition. \**p*=0.0161 and \*\*\**p*= 0.0003, or <0.0001 respectively, two-sided Student’s *t*-test. (D) Quantitative comparison of the timing of CO designation in worms as described in (B). Worms were treated with or without 1 mM auxin for 3, 5 or 7 h before analysis. Quantification was performed as described in (C). \*\*\**p*<0.0001, two-sided Student’s *t*-test.

We reasoned that if CO designation is inhibited by CHK-2, then depletion of CHK-2 should be sufficient to restore earlier designation in mutants that disrupt establishment of CO intermediates on a subset of chromosomes. We depleted CHK-2 in *him-8* and *him-5* mutants, which specifically affect synapsis or DSB induction on the *X* chromosomes, respectively, but do not impair CO formation on autosomes (Hodgkin et al., 1979; Broverman and Meneely, 1994; Phillips et al., 2005; Meneely et al., 2012). In each case we observed a shift in the appearance of bright COSA-1 foci towards the distal region of meiotic prophase following CHK-2 depletion (Figure 2B and D; Figure 2– figure supplement 1C-F). This result reinforces evidence described above that CHK-2 activity is required to delay CO designation in response to feedback regulation.

We observed some differences between the two mutants: in *him-8* mutants RAD-51 foci were far more abundant in the extended region of CHK-2 activity, and persisted even after CHK-2 was depleted, resulting in the appearance of nuclei with both RAD-51 foci and bright COSA-1 foci. In contrast, RAD-51 foci were sparser in *him-5* and mostly disappeared upon CHK-2 depletion (Figure 2–figure supplement 1C-E). These differences likely reflect the different arrest points of the two mutants: *him-8* oocytes are defective in synapsis (Phillips et al., 2005), and thus arrest at a zygotene-like state with high CHK-2 activity, while *him-5* mutants are deficient in DSB initiation on the X chromosome (Meneely et al., 2012), resulting in an prolonged “early-pachytene” state with an intermediate level of CHK-2 activity. We speculate that progression to early pachytene and the associated reduction in CHK-2 activity in *him-*5 mutants may attenuate break formation and/or allow breaks to progress to a more advanced stage of repair, so that upon CHK-2 depletion they are more rapidly resolved.

### Relocalization of Polo-like kinases to SCs promotes CHK-2 inactivation and CO designation

A key unanswered question is how CHK-2 is normally inactivated at mid-prophase to terminate DSB induction and promote CO designation. Prior work on inactivation of DNA damage response (DDR) signaling in proliferating cells revealed that the Polo-like kinase Plk1 enables mammalian cells to enter mitosis following DDR activation by phosphorylating the FHA domain of Chk2, which inhibits substrate binding (Vugt et al., 2010). We wondered whether a similar mechanism might regulate CHK-2 in meiosis.

*C. elegans* expresses multiple homologs of mammalian Plk1, including PLK-1, which is essential for mitosis and thus for viability, and PLK-2, which is dispensable for development but plays important roles in meiosis (Chase et al., 2000; Harper et al., 2011; Labella et al., 2011). Loss of *plk-2* perturbs pairing and synapsis, and also causes a pronounced delay in CO designation and subsequent chromosome remodeling during late prophase (Harper et al., 2011; Labella et al., 2011). To test whether CHK-2 might be a substrate of PLK-2, we performed *in vitro* phosphorylation assays using purified PLK-2. We used purified kinase-dead CHK-2^KD^ as a substrate to avoid potential autophosphorylation of CHK-2. Mass spectrometry identified threonine 120 of CHK-2 as an *in vitro* target of PLK-2 (Figure 3–figure supplement 1A). This highly conserved residue lies within the substrate recognition FHA domain (Figure 3A-B), close to serine 116, which corresponds to a site phosphorylated by Plk1 in human cells during DNA damage checkpoint adaptation (Vugt et al., 2010). S116 and T120 both conform to a consensus motif for Plk1 substrates (Figure 3A) (Santamaria et al., 2011). These results indicated that PLK-2 might phosphorylate and inactivate CHK-2 during meiosis.

**Figure 3.**
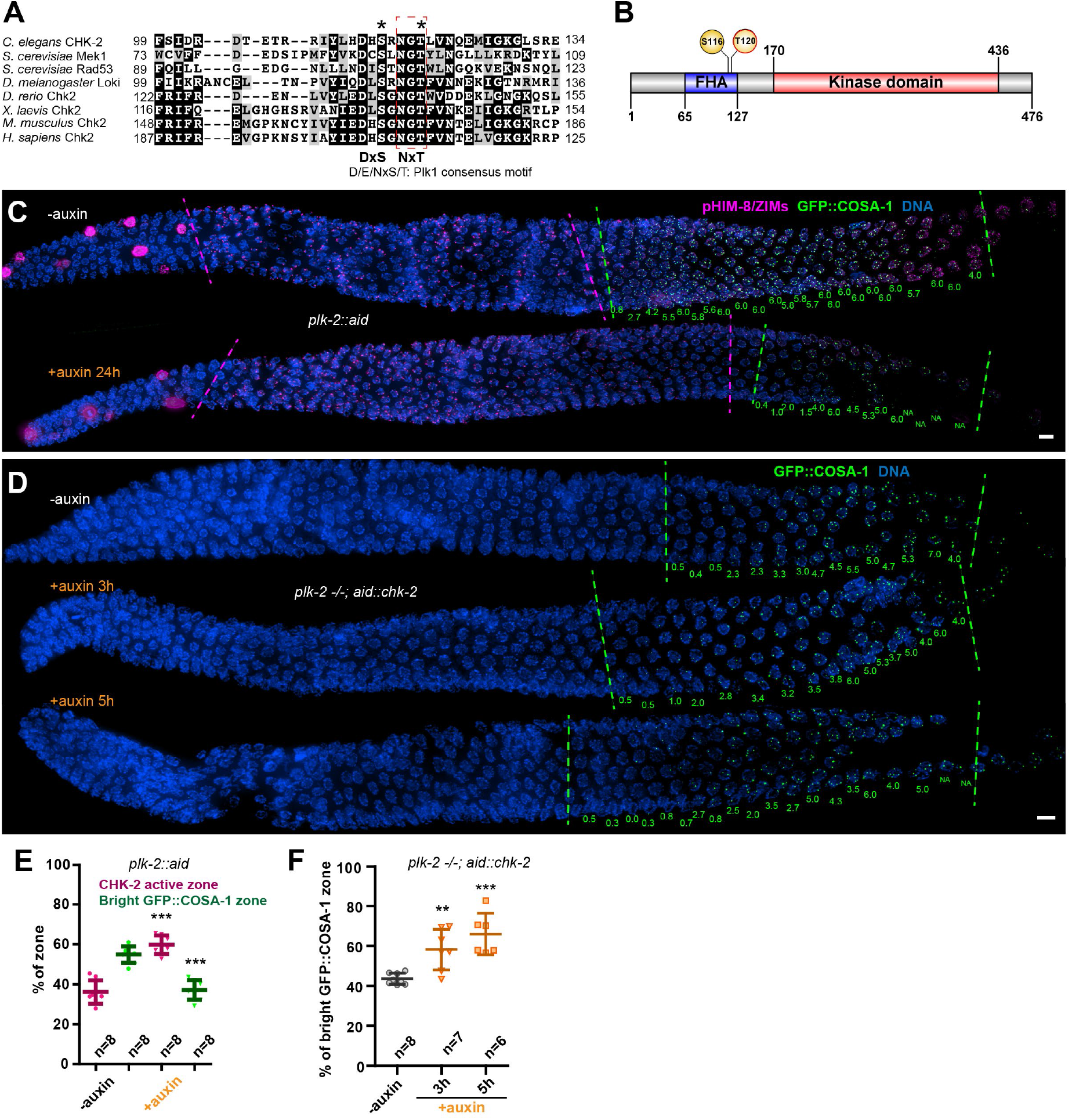
Inactivation of CHK-2 by Polo-like kinase promotes CO designation. (A) Sequence alignment of CHK-2 orthologs from various eukaryotes generated with T-Coffee (Notredame et al., 2000), showing the conservation of Ser116 and Thr120 (asterisks). Thr120 is a direct target of PLK-2; Ser116 corresponds to a Plk1 site identified in mammalian Plk1 (Vugt et al., 2010). Dark and gray shading indicate identical and similar residues, respectively. Both Ser116 and Thr120 match the Plk1 consensus motif [D/E/N]-X-[S/T] (Santamaria et al., 2011). (B) Schematic showing the domain organization of CHK-2 protein and the positions of two phosphorylation sites, Ser116 and Thr120. FHA: Forkhead-associated domain. Numbers indicate amino acid positions. *C. elegans* CHK-2 and budding yeast Mek1 are meiosis-specific kinases that share the FHA and serine/threonine kinase domains of mammalian Chk2 and yeast Rad53, but lack the N-terminal SQ/TQ cluster that regulates activation of Chk2 by ATM. (C) Depletion of PLK-2 delays both CHK-2 inactivation and the appearance of bright COSA-1 foci. Worms were treated with 1 mM auxin for 24 h and stained for pHIM-8/ZIMs (magenta), GFP::COSA-1 (green) and DNA (blue). Dashed magenta lines indicate the CHK-2 active zone. Green lines indicate the bright COSA-1 zone. The average number of bright COSA-1 foci per nuclei in each row is indicated in green below each image. Scale bars, 5 µm. (D) Depletion of CHK-2 restores early appearance of bright COSA-1 foci in *plk*-2 mutants. Worms were treated with or without 1 mM auxin for 3 or 5h. Scale bars, 5 µm. (E-F) Quantification of the extension of the CHK-2 active zone and delay in appearance of bright COSA-1 foci in worms depleted for PLK-2, as described in (C) and of bright COSA-1 foci appearance in *plk-2* mutants upon depletion of CHK-2 as described in (D), respectively. *n* = number of gonads scored for each condition. \*\**p*=0.0018 and \*\*\**p*<0.0001, two-sided Student’s *t*-test.

To determine whether CHK-2 is phosphorylated at Thr120 *in vivo*, we immunoprecipitated epitope-tagged CHK-2 from *C. elegans*. Transcriptome and proteome analyses have found CHK-2 to be a low-abundance protein (WormBase; PAXdb). Consistent with this, tagged CHK-2 was difficult to detect when immunoprecipitated from wild-type animals. Based on other evidence (see below) we suspected that phosphorylation of CHK-2 might lead to its degradation. We thus exploited the AID system to deplete PAS-1, a subunit of the 20S proteasome, as a potential way to increase the abundance of CHK-2. Under conditions in which PAS-1 was efficiently depleted, CHK-2 was clearly enriched (Figure 3–figure supplement 1B-C). Mass spectrometry analyses of the immunoprecipitates revealed that CHK-2 is indeed phosphorylated on Thr120 (Figure 3–figure supplement 1D). To test the role of phosphorylation on the corresponding serine (S116) and/or T120 of CHK-2, we mutated each of these sites to nonphosphorylatable and phosphomimetic residues. Surprisingly, the abundance of all these mutant CHK-2 proteins was much lower than the wild-type protein; only S116A was detectable (Figure 3–figure supplement 2A). Consistently, these mutations all led to loss of CHK-2 function: phopsho-HIM-8/ZIMs was not detectable and bivalent formation was defective; only S116A was partially functional (Figure 3–figure supplement 2B). These indicate that these mutations destabilize CHK-2 in addition to potentially inhibiting substrate binding.

To determine whether PLK-2 is important for meiotic progression independent of its role in homolog pairing and synapsis (Harper et al., 2011; Labella et al., 2011; Sato-Carlton et al., 2017), we used a degron-tagged allele (Zhang et al., 2018). Depletion of PLK-2 significantly delayed CHK-2 inactivation and CO designation (Figure 3C and E). We reasoned that if PLK-2 promotes timely CO designation primarily by inactivating CHK-2, then the delay observed in *plk-2* mutants should be rescued by depletion of CHK-2. Indeed, we found that AID-mediated depletion of CHK-2 restored earlier CO designation in *plk-2* mutants (Figure 3D and F).

As the synaptonemal complex (SC) assembles, PLK-2 is recruited to this structure by binding to a Polo box interacting motif on SYP-1 (Sato-Carlton et al., 2017; Brandt et al., 2020). Following CO designation at mid-pachytene, PLK-2 activity becomes restricted to the “short arms” of each bivalent (Sato-Carlton et al., 2017). We speculated that recruitment of PLK-2 to the SC may promote its ability to inactivate CHK-2. To test this idea, we examined CHK-2 activity in *syp-1*^*T452A*^ mutants, which lack the Polo-box binding motif (S-pT-P) that recruits Polo kinases to the SC (Sato-Carlton et al., 2017; Figure 4A). Consistent with prior analysis, this single point mutation markedly delayed the appearance of bright COSA-1 foci (Sato-Carlton et al., 2017; Figure 4B and D). We found that CHK-2 activity was similarly extended in *syp-1*^*T452A*^ mutants (Figure 4B and D). Depletion of CHK-2 restored earlier CO designation in *syp-1*^*T452A*^ mutants (Figure 4C and E), as in *plk-2* mutants. Thus, binding of PLK-2 to the SC is important for downregulation of CHK-2, and conversely, the delay in CO designation in *syp-1*^*T452A*^ mutants is a direct consequence of persistent CHK-2 activity.

**Figure 4.**
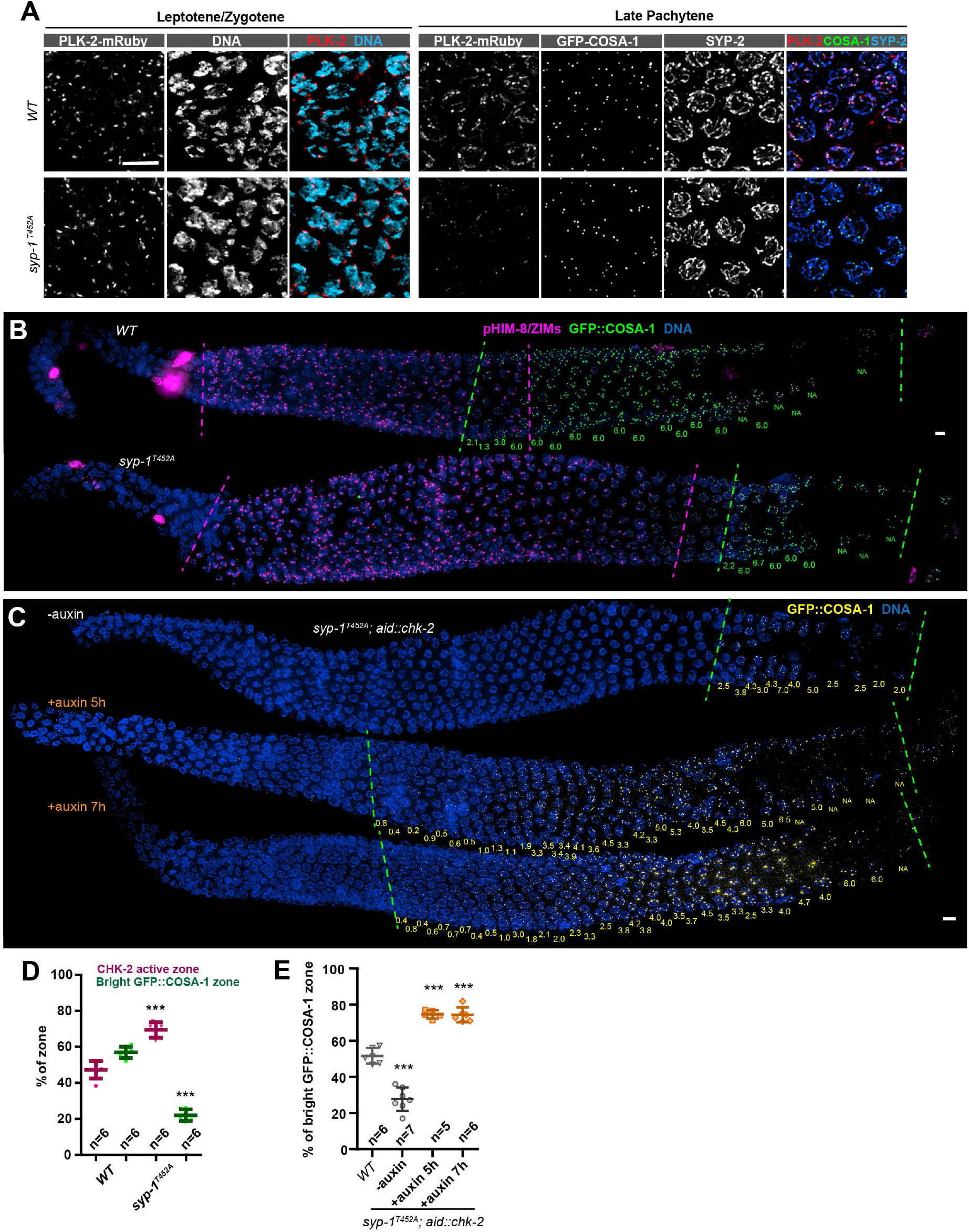
CHK-2 inactivation requires recruitment of Polo-like kinases to the SC. (A) Representative leptotene/zygotene or late pachytene nuclei in wildtype or *syp-1*^*T452A*^ hermaphrodite gonads stained for PLK-2::mRuby (red), DNA (cyan), GFP::COSA-1 (green) and SYP-2 (blue). A conserved Polo box recruitment motif on the SC is absent in *syp-1*^*T452A*^ mutants. PLK-2 localized to pairing centers in leptotene/zygotene nuclei but failed to localize to SC in pachytene nuclei in *syp-1*^*T452A*^ mutants. (B) Representative hermaphrodite gonads stained for pHIM-8/ZIMs (magenta), GFP::COSA-1 (green) and DNA (blue), showing extension of CHK-2 active zone and delayed appearance of bright COSA-1 foci in *syp-1*^*T452A*^ mutants. Dashed magenta lines indicate the CHK-2 active zone, while green lines indicate the bright COSA-1 zone. The average number of bright COSA-1 foci per nuclei in each row are indicated in green below each image. Scale bars, 5 µm. (C) Representative hermaphrodite gonads stained for GFP::COSA-1 (yellow) and DNA (blue), showing that depletion of CHK-2 in *syp-1*^*T452A*^ mutants leads to early appearance of bright COSA-1 foci. Worms were treated with or without 1mM auxin for 5 or 7h. Dash lines indicate the zone with bright COSA-1 foci. Scale bars, 5 µm. (D) Quantification of the extension of CHK-2 active zone and the delay in appearance of bright COSA-1 foci in worms as described in (B). *n* = number of gonads scored for each condition. \*\*\**p*<0.0001, two-sided Student’s *t*-test. (E) Quantification of bright COSA-1 zone in worms maintained and treated as in (C). Wild type worms were used as control. *n* = number of gonads scored for each condition. \*\*\**p*<0.0001, two-sided Student’s *t*-test.

We observed that *plk-2* null mutations, induced depletion of PLK-2, and SYP-1^T452A^ each delayed CO designation to different extents: the relative lengths of bright COSA-1 zone in these strains were 43.61±2.81%, 37.14±4.97% and 20.05±3.24%, respectively. The delay in *plk-2* null mutants was likely affected by other prophase defects, including delayed paring and synapsis (Harper et al., 2011; Labella et al., 2011). Depletion of PLK-2 was less disruptive to these early events (Figure 4–figure supplement 1A). SYP-2^T452A^ caused far milder defects in pairing and synapsis (Brandt et al., 2020; Sato-Carlton et al., 2017), but a more dramatic delay in CHK-2 inactivation and CO designation (Figure 3C-F, 4B and D). This supports the conclusion that recruitment of Polo-like kinase to SCs is important for CHK-2 inactivation and CO designation.

Although mutation or depletion of PLK-2 or failure to recruit it to the SC delayed inactivation of CHK-2, markers for CHK-2 activity eventually disappeared and CO designation was detected in all of these strains. We thus wondered how CHK-2 is inactivated in the absence of PLK-2. In *plk-2* null mutants, PLK-1 can partially substitute at Pairing Centers to promote pairing and synapsis (Harper et al., 2011; Labella et al., 2011). We tested whether PLK-1 might compensate for loss of PLK-2 in late prophase, as it does during early meiosis, by co-depleting both paralogs (Figure 4–figure supplement 1A). Compared to PLK-2 depletion alone, CO designation was further delayed when PLK-1 and PLK-2 were both depleted, indicating that PLK-1 can contribute to CHK-2 inactivation during late prophase (Figure 3C and E; Figure 4–figure supplement 1B and C), but CHK-2 is still inactivated eventually in the absence of both paralogs. We also tested whether the nonessential Plk1 homolog *plk-3* can promote CO designation. As previously reported (Harper et al., 2011), *plk-3* mutants showed no apparent meiotic defects, and we observed no further delay when we combined *plk-3* mutant with depletion of PLK-1 and PLK-2 (data not shown).

The ERK kinase MPK-1 promotes pachytene exit and has been proposed to be important for crossover designation (Church et al., 1995; Hayashi et al., 2007; Lee et al., 2007; Nadarajan et al., 2016). We used the AID system to test whether MPK-1 promotes CHK-2 inactivation or CO designation. MPK-1::AID was undetectable in germ cells following 1 hour of auxin treatment (Figure 4–figure supplement 2A). Although depletion of MPK-1 led to a disordered appearance of the proximal germline, it did not cause any apparent delay in CO designation, either alone or in combination with *syp- 1*^*T452A*^ (Figure 4–figure supplement 2B-E). We thus conclude that MPK-1 activity does not play a role in CO designation, and speculate that a rise in CDK activity and/or other spatially regulated signals in the proximal gonad may lead to CHK-2 inactivation even when Polo-like kinase activity is absent.

## Discussion

In summary, we find that CHK-2 activity is required to inhibit crossover designation. CHK-2 is normally inactivated at mid-pachytene through the recruitment of PLK-2 to the SC and the formation of crossover precursors, but delays in synapsis prevent the activation of PLK-2 and thereby prolong CHK-2 activity. Notably, delays in the establishment of crossover intermediates also prolong CHK-2 activity, and thus full activation of PLK-2 may depend on the formation of crossover intermediates on all chromosomes. Consistent with this idea, PLK-2-dependent phosphorylation of the axis protein HIM-3 (Sato-Carlton et al., 2020) and the central region protein SYP-4 (Nadarajan et al., 2017) increase markedly starting at mid-pachytene, concomitant with CO designation. How the formation of crossover precursors promotes PLK-2 activity remains unclear, but the kinase does appear to associate with designated crossover sites even when it cannot be recruited to the SC (Zhang et al., 2018).

Our data support the idea that PLK-2 directly regulates CHK-2 via inhibitory phosphorylation, but do not rule out the possibility that recruitment of PLK-2 to the SC leads indirectly to CHK-2 inactivation. A recent study reported that CHK-2 inactivation is reversible through mid-pachytene (Castellano-Pozo et al., 2020), consistent with the idea that it occurs through phosphorylation, which can be reversed by phosphatase activity (Kar and Hochwagen, 2021). We speculate that phosphorylated CHK-2 may be degraded after mid-pachytene, resulting in irreversible inactivation.

Intriguingly, both CHK-2 and PLK-2 are active during leptotene/zygotene, when both kinases are bound to Pairing Centers (Kim et al., 2015; Link et al., 2018; Penkner et al., 2009; Sato et al., 2009; Woglar et al., 2013), implying that in this context PLK-2 does not inactivate CHK-2. We speculate that the configuration of CHK-2 and PLK-2 recruitment motifs on the Pairing Center proteins may make the target site(s) in the FHA domain of CHK-2 inaccessible to the active site of PLK-2. However, the activity of CHK-2 in early prophase also depends on an enigmatic meiotic regulator of PLK-2 activity, a heterodimeric complex of HAL-2 and HAL-3 (Roelens et al., 2019; Zhang et al., 2012). This may inhibit or antagonize phosphorylation of CHK-2. Further work will be needed to clarify how CHK-2 and PLK-2 work in concert at pairing centers during early meiosis, while PLK-2 antagonizes CHK-2 later in meiotic prophase.

In *C. elegans*, synapsis can occur normally in the absence of double-strand breaks or recombination intermediates (Dernburg et al., 1998), and meiocytes surveil both synapsis and CO precursors to ensure chiasma formation and faithful segregation. In organisms where synapsis depends on the stabilization of interhomolog joint molecules through the “ZMM” pathway (Pyatnitskaya et al., 2019), cells may monitor synapsis as a proxy for the presence of CO-competent intermediates. However, in either case, SC assembly is essential for the surveillance. In mouse spermatocytes, PLK1 localizes along SCs and promotes pachytene exit (Jordan et al., 2012). Thus, similar mechanisms may coordinate synapsis with meiotic progression in other organisms.

## Materials and methods

### Worm strains

All *C. elegans* strains were maintained on standard nematode growth medium (NGM) plates seeded with OP50 bacteria at 20°C. All epitope-and degron-tagged alleles analyzed in this study were fully functional, as indicating by their ability to support normal meiosis and development (Supplementary Table S1). Unless otherwise indicated, new alleles used in this study were generated by CRISPR/Cas9-mediated genome editing following a modified protocol as previously described (Arribere et al., 2014; Paix et al., 2015; Zhang et al., 2018).

See Supplementary Table S2 for a list of new alleles generated in this study and Supplementary Table S3 for reagents used to make these alleles. A list of worm strains used in this study was shown in Supplementary Table S4. Unless otherwise indicated, young adults (20-24 h post-L4) were used for both immunofluorescence and Western blot assays.

### Worm viability and fertility

To quantify brood sizes, male self-progeny, and embryonic viability, L4 hermaphrodites were picked onto individual seeded plates and transferred to new plates daily over 4 days. Eggs were counted daily. Viable progeny and males were scored when they reached the L4 or adult stages.

### Auxin-mediated protein depletion in worms

Auxin-mediated protein depletion was performed as previously described (Guo et al., 2022; Zhang et al., 2015). Briefly, worms were transferred to bacteria-seeded plates containing 1 mM indole-3-acetic acid (Acros Organics, Cat #122160250) and incubated for the indicated time periods before analysis.

### *In vitro* phosphorylation assay

Recombinant kinase-dead CHK-2 was expressed and purified as described previously (Kim et al., 2015). For PLK-2, the full-length open reading frame (ORF) was amplified from a *C. elegans* cDNA library and cloned into pFastBac1 (Life Technologies) with a GST tag at its N-terminus. GST-PLK-2 was expressed in insect Sf9 cells using the standard Bac-to-Bac system (Life Technologies) and then purified using glutathione Sepharose (GE Life Sciences).

*In vitro* kinase assays were performed in a buffer comprised of 25 mM HEPES pH 7.4, 50 mM NaCl, 2 mM EGTA, 5 mM MgSO4, 1 mM DTT and 0.5 mM NaF, supplemented with 0.5 mM Mg-ATP. 2 µM of GST-CHK-2 KD were incubated with or without 0.2 µM of GST-PLK-2 in the presence or absence of the kinase inhibitor at room temperature for 1 hour. Kinase reactions were terminated by either quick freezing in liquid nitrogen or addition of SDS sample buffer. Proteins in SDS buffer were electrophoresed using gradient polyacrylamide gels (Genscript, #M00652). CHK-2 bands were excised and stored at 4°C. Proteins were then subjected to either in-solution or in-gel trypsin digestion, and phosphorylation sites were identified using mass spectrometry analyses (UC Davis).

### Mass spectrometry

An Xevo G2 QTof coupled to a nanoAcquity UPLC system (Waters, Milford, MA) was used for phosphorylation site identification. Briefly, samples were loaded onto a C18 Waters Trizaic nanotile of 85 um × 100 mm; 1.7 μm (Waters, Milford, MA). The column temperature was set to 45°C with a flow rate of 0.45 mL/min. The mobile phase consisted of A (water containing 0.1% formic acid) and B (acetonitrile containing 0.1% formic acid). A linear gradient elution program was used: 0–40 min, 3–40 % (B); 40-42 min, 40–85 % (B); 42-46 min, 85 % (B); 46-48 min, 85-3 % (B); 48-60 min, 3% (B).

Mass spectrometry data were recorded for 60 minutes for each run and controlled by MassLynx 4.2 (Waters, Milford, MA). Acquisition mode was set to positive polarity under resolution mode. Mass range was set from 50 – 2000 Da. Capillary voltage was 3.5 kV, sampling cone at 25 V, and extraction cone at 2.5 V. Source temperature was held at 110C. Cone gas was set to 25 L/h, nano flow gas at 0.10 Bar, and desolvation gas at 1200 L/h. Leucine–enkephalin at 720 pmol/ul (Waters, Milford, MA) was used as the lock mass ion at *m*/*z* 556.2771 and introduced at 1 uL/min at 45 second intervals with a 3 scan average and mass window of +/-0.5 Da. The MSe data were acquired using two scan functions corresponding to low energy for function 1 and high energy for function 2. Function 1 had collision energy at 6 V and function 2 had a collision energy ramp of 18 − 42 V.

RAW MSe files were processed using Protein Lynx Global Server (PLGS) version 3.0.3 (Waters, Milford, MA). Processing parameters consisted of a low energy threshold set at 200.0 counts, an elevated energy threshold set at 25.0 counts, and an intensity threshold set at 1500 counts. The databank used corresponded to *C. elegans* and was downloaded from uniport.org and then randomized. Searches were performed with trypsin specificity and allowed for two missed cleavages. Possible structure modifications included for consideration were methionine oxidation, carbamidomethylation of cysteine, deamidiation of asparagine or glutamine, dehydration of serine or threonine, and phosphorylation of serine, threonine, or tyrosine.

### ALFA::CHK-2 immunoprecipitation

To obtain synchronized young adults, 5-6 L4 larvae animals were picked onto standard 60-mm culture plate spread with OP50 bacteria. Animals were grown at 20°C for five days until starved. 90 plates of L1 larvae for each genotype or condition were washed into 1.5 L liquid culture supplemented with HB101 bacteria. Worms were then grown with aeration at 200rpm at 20°C for 3 days to reach adulthood. Six hours prior to harvest, 1 mM auxin or 0.25% ethanol (solvent control) was added to the liquid culture. Harvested worms were frozen in liquid nitrogen and stored at -80°C. The frozen worms were then processed using a prechilled Retsch mixer mill to break the cuticle, thawed on ice in cold lysis buffer (25 mM HEPES pH7.4, 100 mM NaCl, 1 mM MgCl_2_, 1 mM EGTA, 0.1% Triton X-100, 1 mM DTT, cOmplete protease inhibitors (Sigma #4693159001) and phosSTOP (Sigma #4906837001)). Lysates were processed using a Dounce homogenizer and sonicated using a Branson Digital Sonifier on ice and then centrifuged at 20,000 RCF for 25 min at 4°C. The supernatant was incubated with anti-ALFA selector (Nanotag Biotechnologies, #N1511) for 3 hours at 4°C. Beads were then washed 6 times with lysis buffer and 3 times with milli-Q water. Proteins on beads were then processed for phosphorylation site identification using mass spectrometry (UC Davis).

Briefly, beads were spun in a 10K MWCO filter (VWR, Radnor, PA) at room temperature for 10 minutes at 10,000 x g and then washed with 50 mM ammonium bicarbonate. Beads were then subjected to reduction at 56°C for 45 minutes in 5.5 mM DTT followed by alkylation for one hour in the dark with iodoacetamide added to a final concentration of 10 mM. The beads were again washed with 50 mM ammonium bicarbonate followed by addition of sequencing grade trypsin to a final enzyme:substrate mass ratio of 1:50 and digested overnight at 37°C. Resultant peptides were then collected in a fresh clean centrifuge tube during a final spin of 16,000xg for 20 minutes. Peptides were dried down in a speed-vac and stored at - 80°C. Prior to analysis, samples were reconstituted in 2% acetonitrile with 0.1% TFA. Samples were then loaded and analyzed as described above.

### Microscopy

Immunofluorescence experiments for *C. elegans* were performed as previously described (Zhang et al., 2018). Images shown in the Figure 4–figure supplement 2A were obtained from worms dissected and fixed in the absence of Tween-20 to retain soluble proteins. Primary antibodies were obtained from commercial sources or have been previously described, and were diluted as follows: Rabbit anti-RAD-51 (1:5,000, Novus Biologicals, #29480002), Rabbit anti-pHIM-8/ZIMs [1:500, (Kim et al., 2015)], Goat anti-SYP-1 [1:300, (Harper et al., 2011), Rabbit anti-SYP-2 [1::500, (Colaiácovo et al., 2003)], Chicken anti-HTP-3 [1:500, (MacQueen et al., 2005)], Mouse anti-HA (1:400, Thermo Fisher, #26183), Mouse anti-GFP (1:500, Millipore Sigma, #11814460001). Mouse anti-FLAG (1:500, Sigma, #F1804), anti-ALFA-At647N (1:500, Nanotag Biotechnologies, N1502-At647N). Secondary antibodies labeled with Alexa 488, Cy3 or Cy5 were purchased from Jackson ImmunoResearch (WestGrove, PA) and used at 1:500. All images were acquired as z-stacks through 8-12 µm depth at z-intervals of 0.2 µm using a DeltaVision Elite microscope (GE) with a 100x, 1.4 N.A. or 60x, 1.42 N.A. oil-immersion objective. Iterative 3D deconvolution, image projection and colorization were carried out using the softWoRx package and Adobe Photoshop CC 2021.

### Western blotting

Adult worms were picked into S-basal buffer and lysed by addition of SDS sample buffer, followed by boiling for 15 min with occasional vortexing. Whole worm lysates were then separated on 4-12% polyacrylamide gradient gels (GenScript, #M00654), transferred to membranes, and blotted with mouse anti-HA (1:1,000, Thermo Fisher, #26183), Guinea pig anti-HTP-3 [1:1,500, (MacQueen et al., 2005)], mouse anti-α-tubulin (1:5,000, Millipore Sigma, #05–829), rabbit anti-β-tubulin (1:2,000, Abcam, ab6046), or HRP-conjugated anti-ALFA sdAb (1:1,000, Nanotag Biotechnologies, N1505-HRP), HRP-conjugated anti-mouse secondary antibodies (Jackson Immunoresearch #115-035-068), HRP-conjugated anti-Rabbit secondary antibodies (Jackson Immunoresearch #111-035-144), HRP-conjugated anti-Guinea pig secondary antibodies (Jackson Immunoresearch #106-035-003). SuperSignal West Femto Maximum Sensitivity Substrate (Thermo Fisher, #34095) were used for detection.

### Timing of CHK-2 inactivation and CO designation

The CHK-2 active zone was determined by immunofluorescence with an antibody recognizing phosphorylated HIM-8 and ZIM proteins (Kim et al., 2015). Designated crossovers were detected using GFP::COSA-1. The lengths of regions corresponding to each meiotic stage vary among individual animals, but the length ratios of these regions are relatively consistent (Kim et al., 2015; Stamper et al., 2013). Thus, unless otherwise indicated, we quantified the CHK-2 active zone or bright COSA-1 zone as a ratio of the region showing positive staining to the total length of the region from meiotic onset to the end of pachytene. Meiotic onset was determined by the staining of meiosis-specific proteins SYP-1 and/or HTP-3.

Bright GFP::COSA-1 foci were identified using the intensity profile of GFP::COSA-1 throughout pachytene, as shown in Figure 1D. While early GFP::COSA-1 foci are relatively dim and sensitive to fixation and antibody staining conditions, COSA-1 foci at designated CO sites are relatively constant in intensity from their appearance until the end of pachytene. The absolute intensity of GFP::COSA-1 varied between samples, so it was not possible to identify designated CO sites using a fixed intensity threshold.

### Quantification and statistical analysis

Quantification methods and statistical parameters are described in the legend of each figure, including sample sizes, error calculations (SD or SEM), statistical tests, and *p*-values. *p*<0.05 was considered to be significant.

## Acknowledgements

This work was supported by funding from the National Institutes of Health (R01 GM065591) and an Investigator award from the Howard Hughes Medical Institute to A.F.D. Some strains were provided by the Caenorhabditis Genetics Center (CGC), which is funded by NIH Office of Research Infrastructure Programs (P40 OD010440). We thank Yumi Kim for sharing the CHK-2 and PLK-2 plasmids, Andrew Ziesel, Nancy Hollingsworth, and members of the Dernburg lab for helpful discussions throughout this work.

## Author contributions

Liangyu Zhang: conceptualization, data curation, formal analysis, investigation, methodology, project administration, resources, software, supervision, validation, visualization, and writing (original draft and review and editing).

Weston T. Stauffer: investigation, methodology and writing (review and editing).

John S. Wang: investigation, methodology, software, validation, and writing (review and editing).

Fan Wu: investigation, methodology, and writing (review and editing).

Zhouliang Yu: investigation, methodology, and writing (review and editing).

Chenshu Liu: investigation, methodology, and writing (review and editing).

Abby F. Dernburg: conceptualization, funding acquisition, project administration, supervision, visualization, and writing (original draft and review and editing).

**Figure 2—figure supplement 1.**
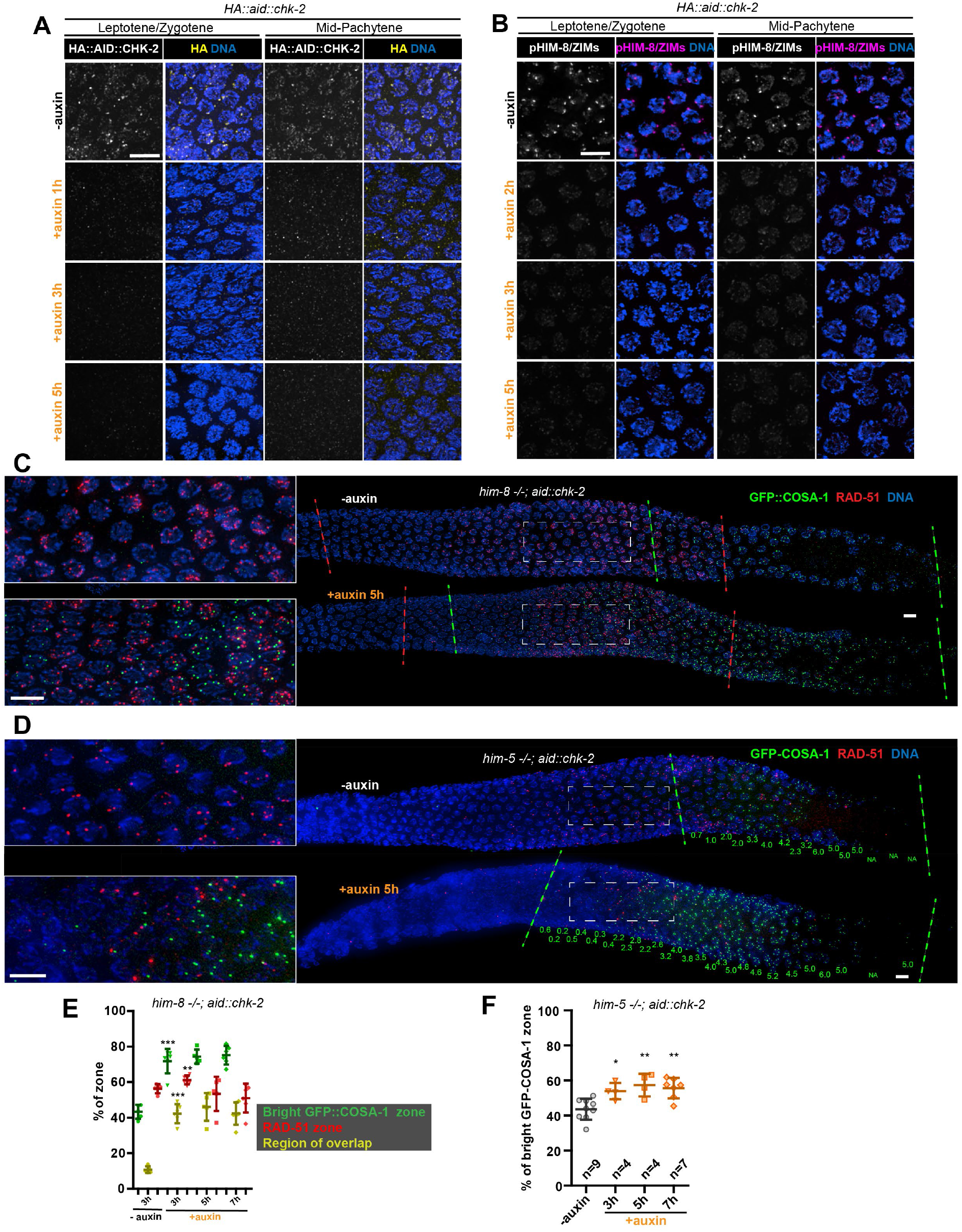
Depletion of CHK-2 activity results in earlier crossover designation. (A) Representative leptotene/zygotene and mid-pachytene nuclei stained for HA::AID::CHK-2 (anti-HA) and DNA, showing efficient depletion of CHK-2 following auxin treatment. Worms were incubated with 1 mM auxin (or 0.25% ethanol lacking auxin) for 1, 3 or 5 h before analysis. CHK-2 was undetectable within 3h of auxin exposure. Scale bars, 5 µm. (B) Representative leptotene/zygotene and mid-pachytene nuclei stained for phosphorylated HIM-8/ZIMs and DNA, showing efficient depletion of CHK-2 activity within 3 h of auxin treatment. Scale bars, 5 µm. (C-D) Representative germlines from *him-8* (C) and *him-5* (D) mutant hermaphrodites stained for GFP::COSA-1 (green), RAD-51 (red) and DNA (blue), revealing earlier appearance of bright COSA-1 foci upon CHK-2 depletion. Worms were transferred to plates prepared with 1 mM auxin (or 0.25% ethanol as a control) for 5 h before fixation. Dashed green lines indicate where bright COSA-1 foci start to appear and where pachytene ends. Dashed red lines in (C) indicate RAD-51-positive zone. Note: the same image in (C) without RAD-51 is shown in Figure 2B. The average number of bright COSA-1 foci per nucleus in each row is indicated below each image in green in (D). Scale bars, 5 µm. (E-F) Quantitative comparison of the RAD-51-positive zone upon CHK-2 depletion in *him-8* mutants and the timing of appearance of bright COSA-1 foci upon CHK-2 depletion in *him-5* mutants. Worms were exposed to auxin for 3, 5 or 7 h before fixation and analysis. The bright GFP::COSA-1 zone was defined as a region from where bright COSA-1 foci start to appear to the end of pachytene, before oocytes form a single row. The “% of bright GFP::COSA-1 zone” in the graph was defined as the ratio of the length of bright GFP::COSA-1 zone to the length of the region from meiotic onset to the end of pachytene. Meiotic onset was determined by the localization of meiosis-specific proteins SYP-1 and/or HTP-3. The RAD-51 zone and the region of overlap between the RAD-51 and bright COSA-1 zones were measured and expressed in the same way. See “materials and methods” for details. \*\**p*=0.001 and \*\*\**p*<0.0001 in (E), \**p*=0.0114, \*\**p*=0.0032 and 0.0012 in (F), respectively, two-sided Student’s *t*-test.

**Figure 3—figure supplement 1.**
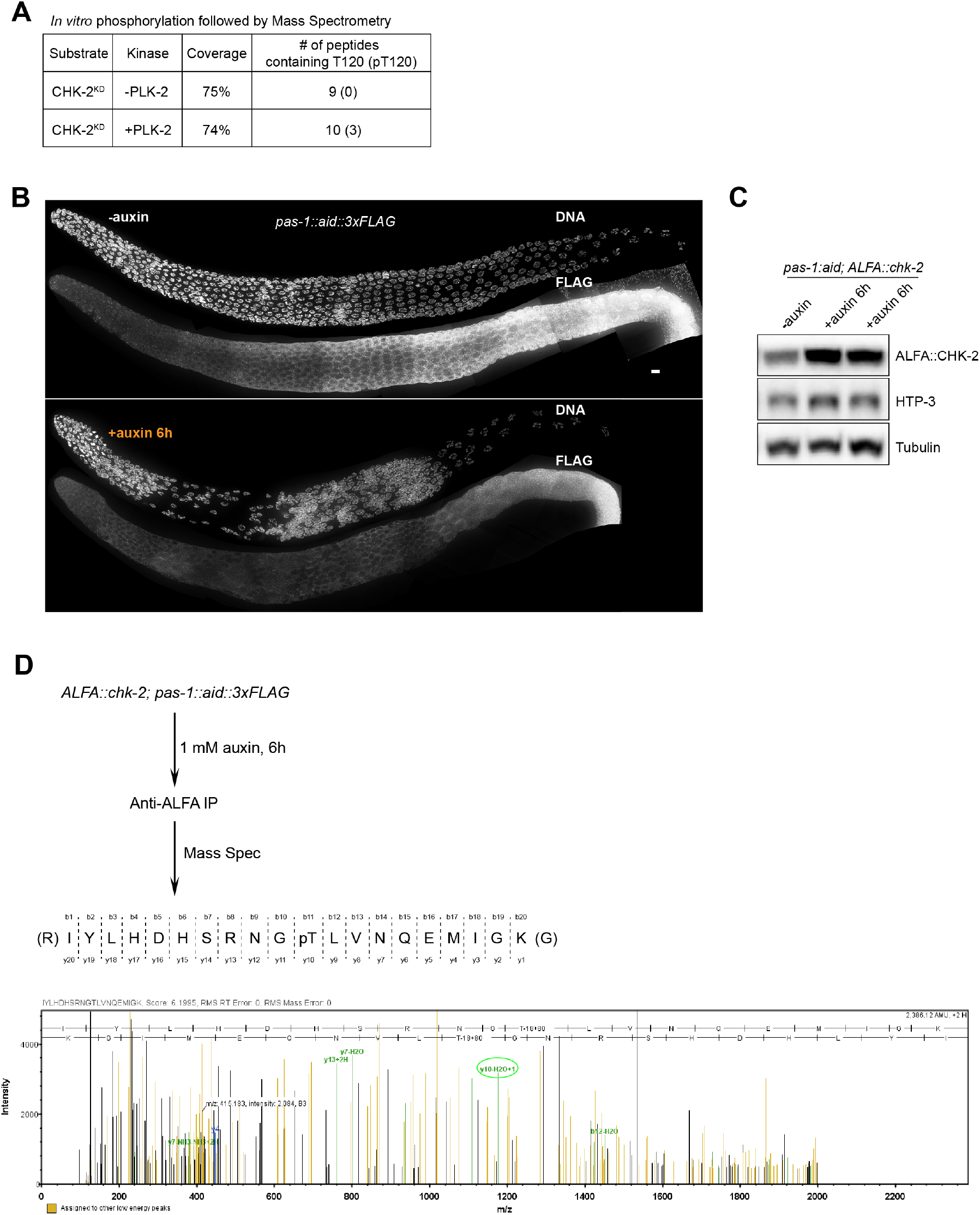
Phosphorylation of CHK-2 by PLK-2. (A) Recombinant kinase-dead CHK-2 protein (CHK-2^KD^) was phosphorylated *in vitro* by PLK-2 kinase and then subjected to mass spectrometry analysis for phosphorylation site identification. CHK-2 Thr120 was identified as a target of PLK-2. (B) Representative hermaphrodite gonads stained for PAS-1::AID::3xFLAG (anti-3xFLAG) and DNA, showing efficient depletion of PAS-1 in the germline upon 1mM auxin treatment for 6 h. PAS-1 is a subunit of the 20S proteasome and is essential for the proteasome activity. Scale bars, 5 µm. (C) A western blot shows increased abundance of ALFA::CHK-2 following depletion of proteasome subunit PAS-1. Worms were transferred to media containing 1 mM auxin for 6 h before analysis. ALFA::CHK-2 was blotted using anti-ALFA nanobody. HTP-3 and tubulin were blotted as loading controls. The two right lanes are biological replicates. (D) ALFA::CHK-2 was immunoprecipitated from adult hermaphrodites following depletion of PAS-1 for 6 hours. Mass spectrometry identified CHK-2 Thr120 as a phosphorylated site *in vivo*.

**Figure 3—figure supplement 2.**
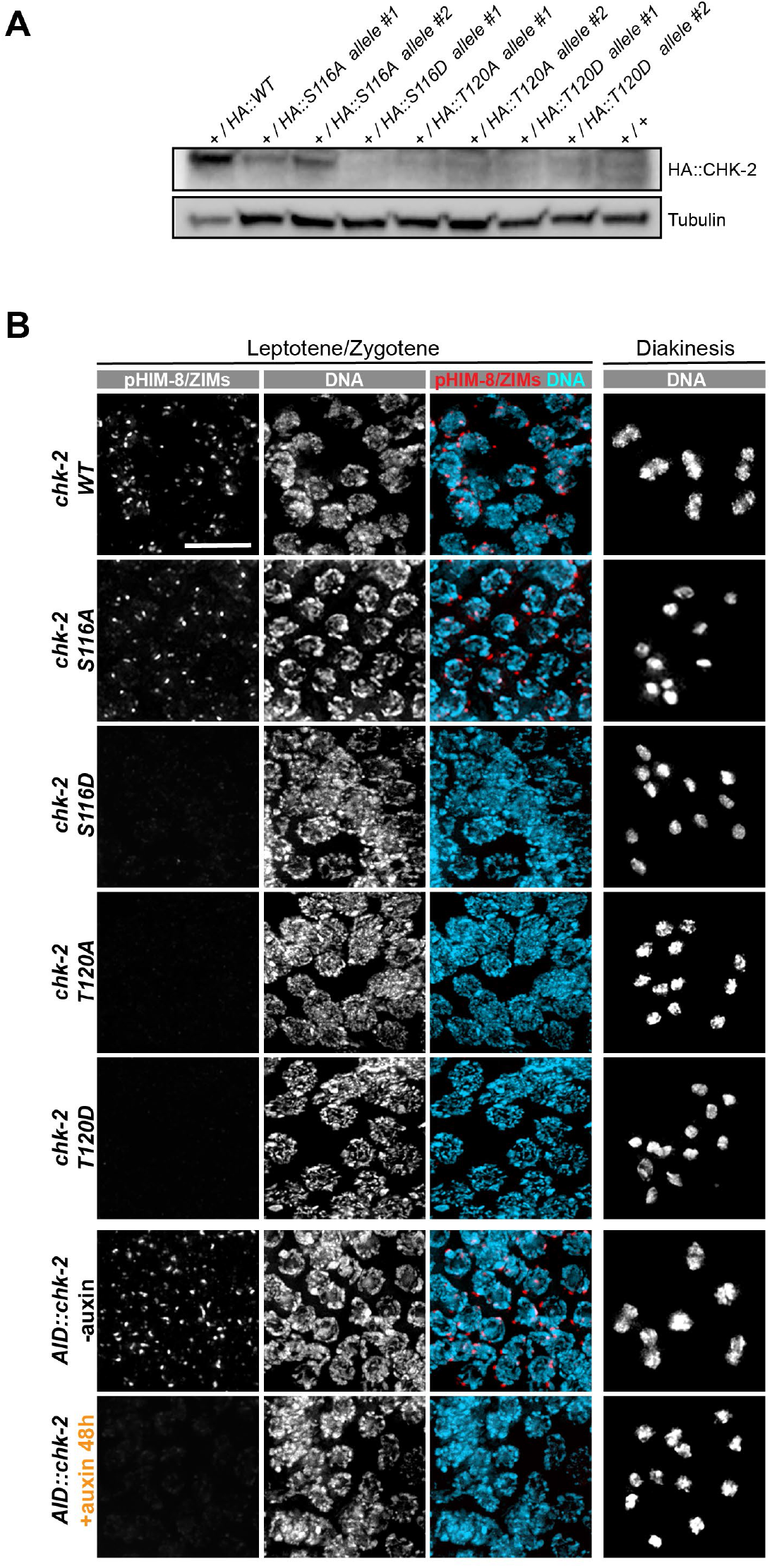
Characterization of CHK-2 phospho-mutants. (A) Worm strains expressing the indicated HA-tagged wild-type and mutant CHK-2 proteins were compared by western blot: HA::CHK-2 WT, S116A, S116D, T120A, and T120D were all measured in heterozygotes as these point mutations result in very few viable progeny and cannot be readily balanced. HA::CHK-2 WT and S116A were readily detected, while other mutant proteins were near or below the limit of detection. HA::CHK-2 was blotted using anti-HA antibody. Tubulin was probed as a loading control. (B) Representative leptotene/zygotene and diakinesis nuclei stained for phosphorylated HIM-8/ZIMs and DNA, showing CHK-2 activity and chiasma formation in CHK-2 WT, S116A, S116D, T120A, T120D, and CHK-2 depleted homozygotes. For auxin treatment, worms were incubated with 1 mM auxin (or 0.25% ethanol lacking auxin) for 48 h before analysis. Like CHK-2 depleted worms, CHK-2 activity and bivalents were not detected in CHK-2 S116D, T120A and T120D mutants. Scale bars, 5 µm.

**Figure 4—figure supplement 1.**
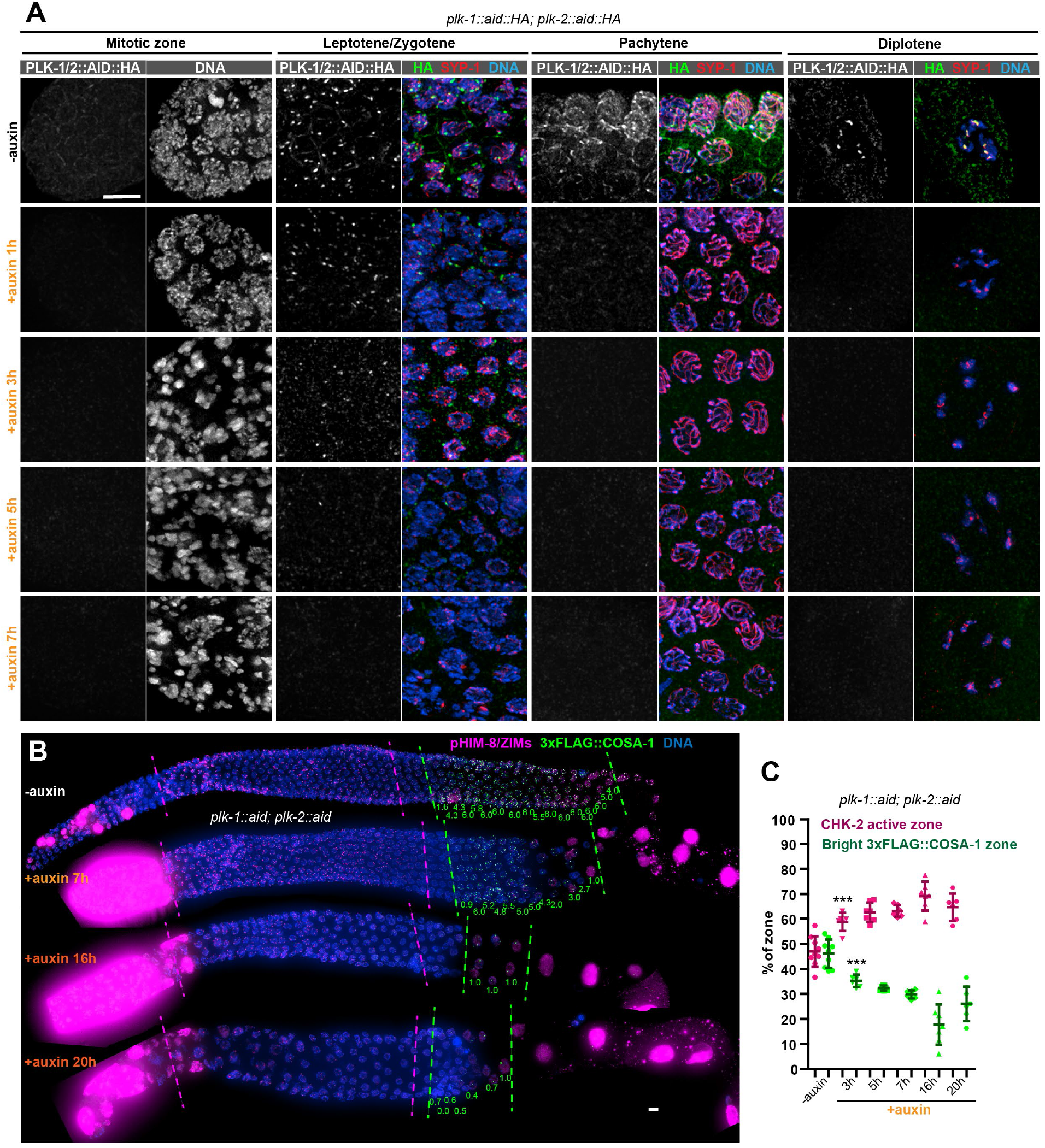
PLK-1 is not required to override the crossover assurance checkpoint in the absence of PLK-2. (A) Representative nuclei at different stages showing efficient co-depletion of PLK-1 and PLK-2 in the germline. One hour of auxin treatment was sufficient to eliminate detectable PLK-1/PLK2 in the germline except at Pairing Centers in early meiotic nuclei, where full depletion required 5 h. Scale bars, 5 µm. (B) Images of representative hermaphrodite gonads stained for pHIM-8/ZIMs (magenta), 3xFLAG-COSA-1 (green) and DNA (blue), showing inactivation of CHK-2 and appearance of bright COSA-1 foci in the absence of PLK-1 and PLK-2. Dashed magenta lines indicate CHK-2 active zone, while green lines indicate the region of nuclei with bright COSA-1 foci. When PLK-1 and PLK-2 were co-depleted for longer than 16 hours, COSA-1 foci were greatly diminished (in most nuclei, only 1 bright 3xFLAG::COSA-1 focus can be detected). This likely reflects defects in pairing and synapsis, which prevent the formation of crossover precursors. However, bright COSA-1 foci were still observed when both PLK-1 and PLK-2 were depleted. Since COSA-1 appear only after inactivation of crossover assurance checkpoint, this suggested that crossover assurance checkpoint can still be overridden in the absence of PLK-1 and PLK-2. The average number of bright COSA-1 foci per nucleus in each row was indicated below each image in green. Scale bars, 5 µm. (C) Graphs showing extension of CHK-2 active zone and delay in appearance of bright COSA-1 foci in worms co-depleted for PLK-1 and PLK-2. The bright COSA-1 zones and CHK-2-active zones were measured and presented as described in Figure 2–figure supplement 1E. *n* = 9, 6, 7, 6, 7 and 6 gonads respectively. \*\*\**p*=0.0007 and 0.0009, respectively, two-sided Student’s *t*-test.

**Figure 4—figure supplement 2.**
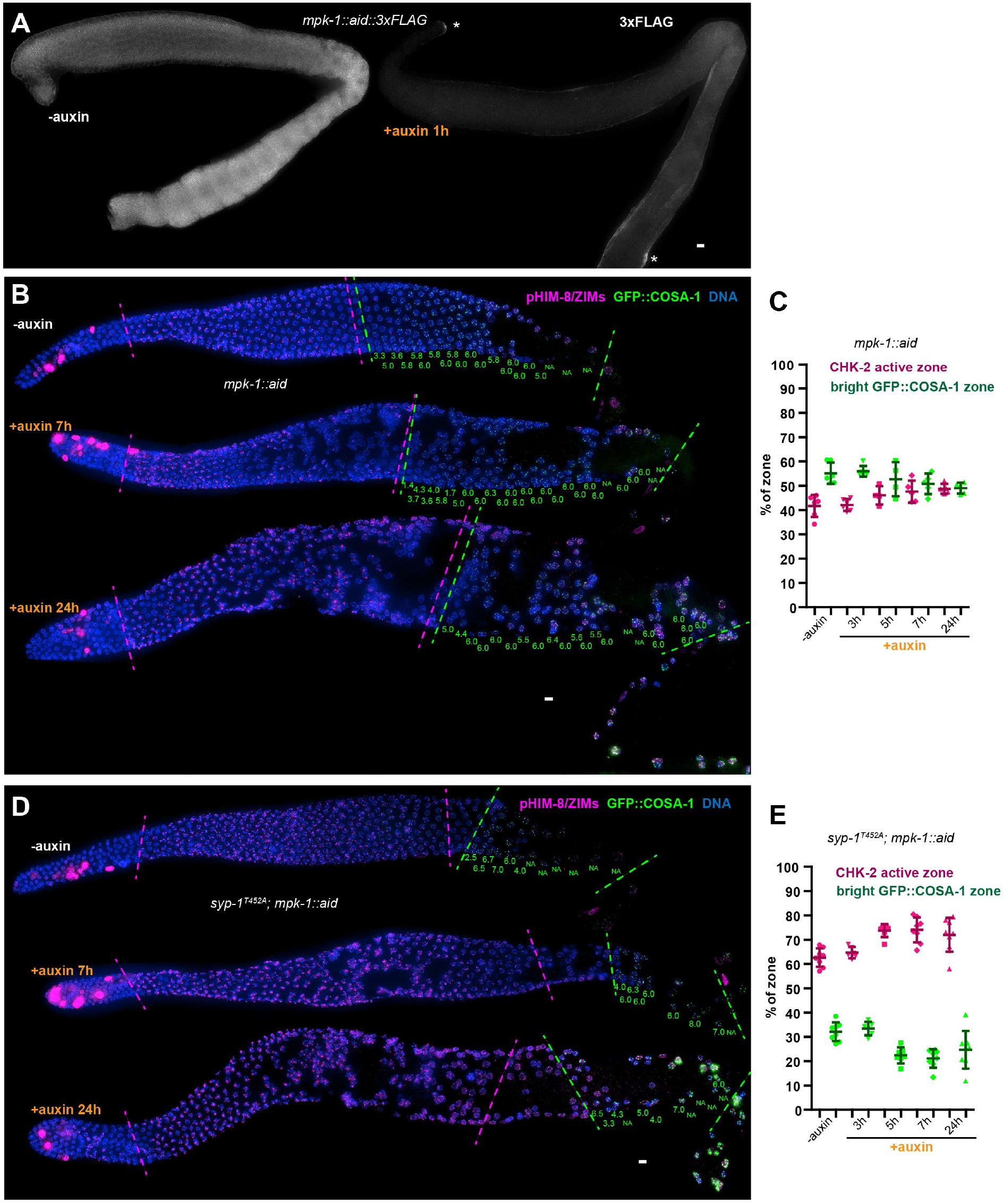
MPK-1 does not silence the crossover assurance checkpoint. (A) Representative hermaphrodite gonads showing efficient, germline-specific depletion of MPK-1. Asterisks indicate the somatic distal tip and seam cells, where MPK-1 persists following degradation since TIR1 is expressed only in germ cells. Scale bars, 5 µm. (B) Representative hermaphrodite gonads stained for pHIM-8/ZIMs (magenta), GFP::COSA-1 (green) and DNA (blue), showing inactivation of CHK-2 and appearance of bright COSA-1 foci in the absence of MPK-1. Magenta dashed lines indicate CHK-2 active zone, while green lines indicate bight COSA-1 zone. The number of bright COSA-1 foci per nucleus in each row was indicated at the bottom of each image in green. Scale bars, 5 µm. (C) Graphs showing CHK-2 active zone and bright COSA-1 zone in worms depleted for MPK-1 for various length of time. In this case, the bright GFP::COSA-1 zones and CHK-2-active zones were measured and presented as a fraction of the region from meiotic onset to the turn of the gonad instead of the end of pachytene, because MPK-1 depletion perturbs pachytene exit and formation of a single line of oocytes. *n* = 6, 6, 4, 5 and 5 gonads respectively. (D) Images of representative hermaphrodite gonads stained for pHIM-8/ZIMs (magenta), GFP::COSA-1 (green) and DNA (blue), showing inactivation of CHK-2 and bright COSA-1 foci formation in the *syp-1*^*T452A*^ worms depleted for MPK-1. Scale bars, 5 µm. (E) Quantification of the CHK-2 active zone and bright COSA-1 zone in *syp-1*^*T452A*^ hermaphrodites depleted for MPK-1 for various lengths of time. ‘bright GFP::COSA-1 zone’ and ‘CHK-2 active zone’ were measured and presented as described in (C). *n* = 8, 6, 7, 8 and 8 gonads respectively.

## Additional files

### Source Data

Figure 3–figure supplement 1–Source Data 1 includes western blotting raw images in Figure 3–figure supplement 1.

Figure 3–figure supplement 2–Source Data 1 includes western blotting raw images in Figure 3–figure supplement 2.

## Supplementary file

Supplementary file 1. This file includes four tables (Table S1-S4). Table S1 reports the viability and fertility of representative transgenic worm strains used in this study, which indicates that all epitope-and degron-tagged alleles support normal meiosis and development. Table S2 lists the worm alleles generated in this study. Table S3 lists the crRNA, repair templates and genotyping primers generated in this study. Table S4 lists the worm strains used in this study.

---

**Supplementary Table S1.** Viability and fertility of transgenic worm strains

Viability and fertility of transgenic worm strains
| Strains | Eggs laid (±SD)<br>(n=6 broods) | Egg viability<br>(±SD) (%) | Male progeny<br>(±SD) (%) |
| --- | --- | --- | --- |
| WT | 261(±17) | 106.64(±5.07) | 0.13(±0.30) |
| *mels8[pie-1p::GFP::cosa-1, unc-119(+)] II*;<br>*ieSi38[sun-1p::tir1::mRuby::sun-1 3' UTR, Cbr-unc-119 (+)] IV*; *chk-2(ie121[HA::aid::chk-2]) V* | 295(±26.51) | 105.63(±3.09) | 0.27(±0.25) |
| *mels8[pie-1p::GFP::cosa-1, unc-119(+)] II*;<br>*ieSi38[sun-1p::tir1::mRuby::sun-1 3' UTR, Cbr-unc-119 (+)] IV*; *chk-2(ie127[HA::chk-2]) V* | 202.33(±24.12) | 103.68(±3.64) | 0.15(±0.38) |
| *plk-2(ie50[plk-2::mRuby]; mels8[pie-1p::GFP::cosa-1, unc-119(+)] II*; *mpk-1(ie54[mpk-1:: aid::3xFLAG]) III*;<br>*ieSi38[sun-1p::tir1::mRuby::sun-1 3' UTR, Cbr-unc-119 (+)] IV* | 274.6(±24.09) | 104.07(±3.11) | 0.15(±0.20) |
| *ieSi64[gld-1p::tir1::mRuby::gld-1 3' UTR, Cbr-unc-119 (+)] II* | 269(±42.53) | 104.41(±1.88) | 0.07(±0.18) |
| *plk-2(ie126[plk-2::aid::HA]) I*; *ieSi64[gld-1p::tir1::mRuby::gld-1 3' UTR, Cbr-unc-119 (+)] II*;<br>*cosa-1(ie124[3xFLAG::cosa-1], plk-1(ie125[plk-1::aid::HA]) III* | 300.83(±40.66) | 101.02(±1.61) | 0.00(±0.00) |
| *mels8[pie-1p::GFP::cosa-1, unc-119(+)] II*;<br>*ieSi38[sun-1p::tir1::mRuby::sun-1 3' UTR, Cbr-unc-119 (+)] IV*; *pas-1(ie185[pas-1::aid::3xFLAG])*, *chk-2(ie186 [ALFA::chk-2]) V* | 217.50(±33.75) | 103.57(±4.26) | 0.05(±0.14) |
| *plk-2(ie126[plk-2::aid::HA]) I*; *ieSi64[gld-1p::tir1::mRuby::gld-1 3' UTR, Cbr-unc-119 (+)] II*;<br>*cosa-1(ie124[3xFLAG::cosa-1], plk-1(ie125[plk-1::aid::HA]) III*; *pas-1(ie188[pas-1::aid::3xFLAG])*, *chk-2(ie187 [ALFA::chk-2]) V* | 245.50(±14.10) | 106.71(±5.17) | 0.00(±0.00) |

Note: viability >100% is a consequence of failing to count some embryos.

**Supplementary Table S2.** Worm alleles generated in this study

Worm alleles generated in this study
| Allele | Genotype | Information about mutagenesis |
| --- | --- | --- |
| ie121 | *chk-2*(ie121[HA::aid::chk-2]) | generated using *dpy-10* Co-CRISPR in *ieSi38*[*sun-1p::tir1::mRuby::sun-1 3'UTR, Cbr-unc-119(+)*] IV |
| ie122 | *him-8* (ie122) mutant | generated using *dpy-10* Co-CRISPR in *mels8*[*pie-1p::GFP::cosa-1, unc-119(+)*] II; *ieSi38*[*sun-1p::tir1::mRuby::sun-1 3'UTR, Cbr-unc-119(+)*] IV; *chk-2*(ie121[HA::aid::chk-2]) V |
| ie123 | *syp-1*(ie123[*syp-1 T452A*]) | generated using *dpy-10* Co-CRISPR in *plk-2*(ie50[*plk-2::mRuby*] I; *mels8*[*pie-1p::GFP::cosa-1, unc-119(+)*] II |
| ieSi64 | [*gld-1p::tir1::mRuby::gld-1 3' UTR, Cbr-unc-119 (+)*] II | Single copy transgene inserted into Chr II (oxTi179) using mosSCI |
| ie124 | *cosa-1*(ie124[3xFLAG::cosa-1]) | generated using *dpy-10* Co-CRISPR in *ieSi64*[*gld-1p::tir1::mRuby::sun-1 3' UTR, Cbr-unc-119 (+)*] II |
| ie125 | *plk-1*(ie125[*plk-1::aid::HA*]) | generated using *dpy-10* Co-CRISPR in *ieSi64*[*gld-1p::tir1::mRuby::sun-1 3' UTR, Cbr-unc-119 (+)*] II; *cosa-1*(ie124[3xFLAG::cosa-1]) III |
| ie126 | *plk-2*(ie126[*plk-2::aid::HA*]) | generated using *dpy-10* Co-CRISPR in *ieSi64*[*gld-1p::tir1::mRuby::sun-1 3' UTR, Cbr-unc-119 (+)*] II; *cosa-1*(ie124[3xFLAG::cosa-1]) III |
| ie54 | *mpk-1*(ie54[*mpk-1::aid::3xFLAG*]) | generated using *dpy-10* Co-CRISPR in *plk-2*(ie50[*plk-2::mRuby*] I; *mels8*[*pie-1p::GFP::cosa-1, unc-119(+)*] II; *ieSi38*[*sun-1p::tir1::mRuby::sun-1 3' UTR, Cbr-unc-119 (+)*] IV |
| ie127 | *chk-2*(ie127[HA::chk-2]) | generated using *dpy-10* Co-CRISPR in *mels8*[*pie-1p::GFP::cosa-1, unc-119(+)*] II; *ieSi38*[*sun-1p::tir1::mRuby::sun-1 3' UTR, Cbr-unc-119 (+)*] IV |
| ie128 | *chk-2*(ie128[HA::chk-2 S116A]) | generated using *dpy-10* Co-CRISPR in *mels8*[*pie-1p::GFP::cosa-1, unc-119(+)*] II; *ieSi38*[*sun-1p::tir1::mRuby::sun-1 3' UTR, Cbr-unc-119 (+)*] IV; *chk-2*(ie127[HA::chk-2]) |
| ie129 | *chk-2*(ie129[HA::chk-2 S116A]) | generated using *dpy-10* Co-CRISPR in *mels8*[*pie-1p::GFP::cosa-1, unc-119(+)*] II; *ieSi38*[*sun-1p::tir1::mRuby::sun-1 3' UTR, Cbr-unc-119 (+)*] IV; *chk-2*(ie127[HA::chk-2]) |
| ie130 | *chk-2*(ie130[HA::chk-2 S116D]) | generated using *dpy-10* Co-CRISPR in *mels8*[*pie-1p::GFP::cosa-1, unc-119(+)*] II; *ieSi38*[*sun-1p::tir1::mRuby::sun-1 3' UTR, Cbr-unc-119 (+)*] IV; *chk-2*(ie127[HA::chk-2]) |
| ie131 | *chk-2*(ie131[HA::chk-2 T120A]) | generated using *dpy-10* Co-CRISPR in *mels8*[*pie-1p::GFP::cosa-1, unc-119(+)*] II; *ieSi38*[*sun-1p::tir1::mRuby::sun-1 3' UTR, Cbr-unc-119 (+)*] IV; *chk-2*(ie127[HA::chk-2]) |
| ie132 | *chk-2*(ie132[HA::chk-2 T120A]) | generated using *dpy-10* Co-CRISPR in *mels8*[*pie-1p::GFP::cosa-1, unc-119(+)*] II; *ieSi38*[*sun-1p::tir1::mRuby::sun-1 3' UTR, Cbr-unc-119 (+)*] IV; *chk-2*(ie127[HA::chk-2]) |
| ie133 | *chk-2*(ie133[HA::chk-2 T120D]) | generated using *dpy-10* Co-CRISPR in *mels8*[*pie-1p::GFP::cosa-1, unc-119(+)*] II; *ieSi38*[*sun-1p::tir1::mRuby::sun-1 3' UTR, Cbr-unc-119 (+)*] IV; *chk-2*(ie127[HA::chk-2]) |
| ie134 | *chk-2*(ie134[HA::chk-2 T120D]) | generated using *dpy-10* Co-CRISPR in *mels8*[*pie-1p::GFP::cosa-1, unc-119(+)*] II; *ieSi38*[*sun-1p::tir1::mRuby::sun-1 3' UTR, Cbr-unc-119 (+)*] IV; *chk-2*(ie127[HA::chk-2]) |
| ie135 | *plk-3*(ie135) mutant | generated using *dpy-10* Co-CRISPR in *mels8*[*pie-1p::GFP::cosa-1, unc-119(+)*] II; *ieSi38*[*sun-1p::tir1::mRuby::sun-1 3' UTR, Cbr-unc-119 (+)*], *him-8*(ie122) IV; *chk-2*(ie121[HA::aid::chk-2]) V |
| ie136 | *plk-3*(ie136) mutant | generated using *dpy-10* Co-CRISPR in *plk-2*(ie126[*plk-2::aid::HA*] I; *ieSi64*[*gld-1p::tir1::mRuby::sun-1 3' UTR, Cbr-unc-119 (+)*] II; *cosa-1*(ie124[3xFLAG::cosa-1], *plk-1*(ie125[*plk-1::aid::HA*]) III |
| ie185 | *pas-1*(ie185[*pas-1::aid::3xFLAG*]) | generated using *dpy-10* Co-CRISPR in *mels8*[*pie-1p::GFP::cosa-1, unc-119(+)*] II; *ieSi38*[*sun-1p::tir1::mRuby::sun-1 3' UTR, Cbr-unc-119 (+)*] IV |
| ie186 | chk-2(ie186[ALFA::chk-2]) | generated using dpy-10 Co-CRISPR in mels8[pie-1p::GFP::cosa-1, unc-119(+)] II; ieSi38[sun-1p::tir1::mRuby::sun-1 3' UTR, Cbr-unc-119 (+)] IV; pas-1(ie185[pas-1::aid::3xFLAG]) V |
| ie187 | chk-2(ie187[ALFA::chk-2]) | generated using dpy-10 Co-CRISPR in plk-2(ie126[plk-2::aid::HA]) I; ieSi64[gld-1p::tir1::mRuby::gld-1 3' UTR, Cbr-unc-119 (+)] II; cosa-1(ie124[3xFLAG::cosa-1], plk-1(ie125[plk-1::aid::HA]) III |
| ie188 | pas-1(ie188[pas-1::aid::3xFLAG]) | generated using dpy-10 Co-CRISPR in plk-2(ie126[plk-2::aid::HA]) I; ieSi64[gld-1p::tir1::mRuby::gld-1 3' UTR, Cbr-unc-119 (+)] II; cosa-1(ie124[3xFLAG::cosa-1], plk-1(ie125[plk-1::aid::HA]) III; chk-2(ie187[ALFA::chk-2]) V |

**Supplementary Table S3.**
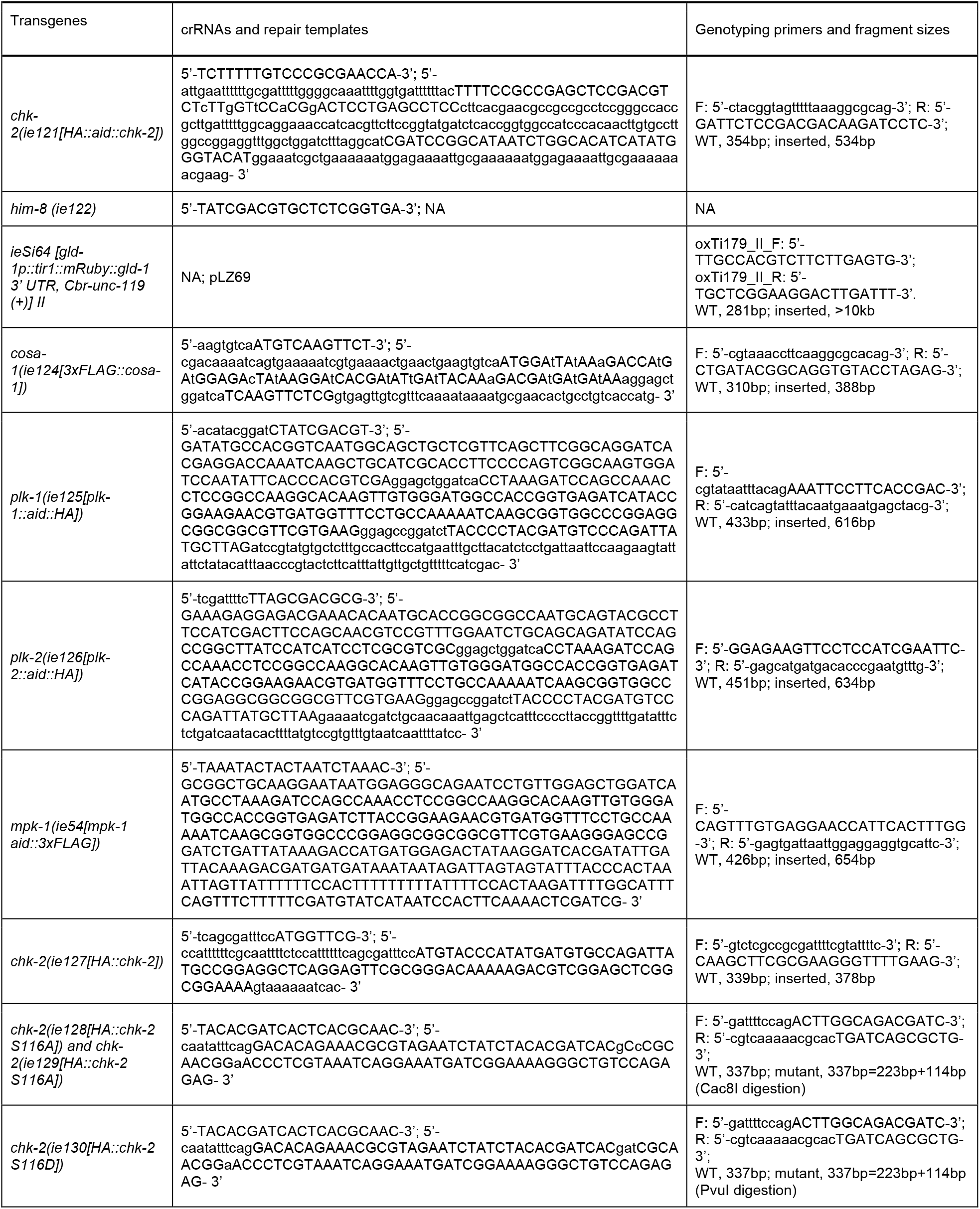
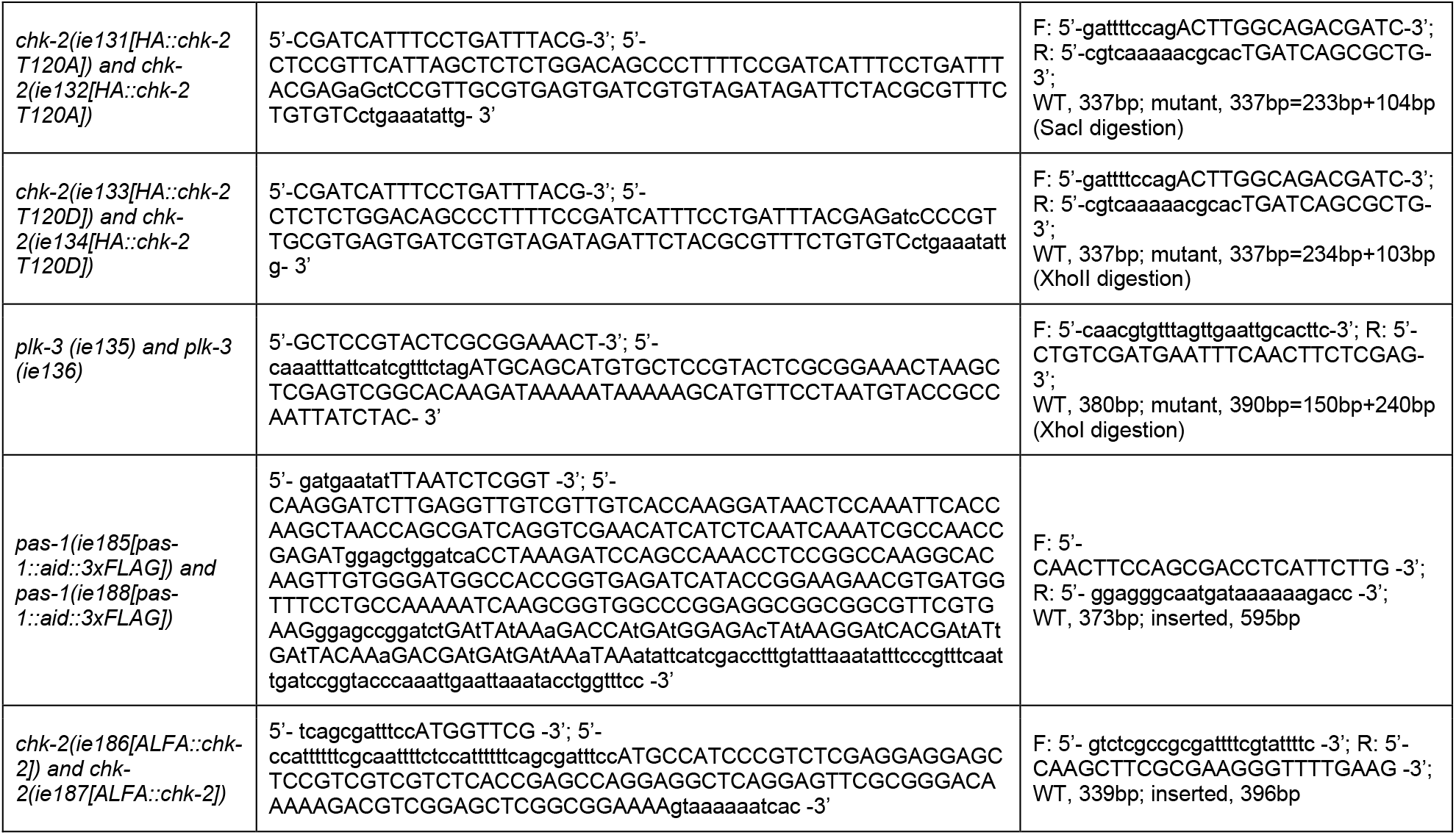
crRNAs, repair templates and genotyping primers used in this study

**Supplementary Table S4.** Worm strains used in this study

Worm strains used in this study
| Strains | Source | Identifier |
| --- | --- | --- |
| *C. elegans*: N2 Bristol, wild isolate | Caenorhabditis Genetics Center | N2 |
| *C. elegans*: *plk-2*(*ie50*[*plk-2*::*mRuby*] I; *mels8*[*pie-1p*::GFP::*cosa-1*, *unc-119*(+)] II; *syp-1*(*ie123*[*syp-1* T452A]) V | This paper | CA1290 |
| *C. elegans*: *ieSi64*[*gld-1p*::*tir1*::*mRuby*::*gld-1* 3' UTR, *Cbr-unc-119* (+)] II | This paper | CA1352 |
| *C. elegans*: *mels8*[*pie-1p*::GFP::*cosa-1*, *unc-119*(+)] II; *ieSi38*[*sun-1p*::*tir1*::*mRuby*::*sun-1* 3' UTR, *Cbr-unc-119* (+)] IV | This paper | CA1364 |
| *C. elegans*: *plk-2*(*ie50*[*plk-2*::*mRuby*] I; *mels8*[*pie-1p*::GFP::*cosa-1*, *unc-119*(+)] II; *mpk-1*(*ie54*[*mpk-1*::*aid*::3xFLAG]) III; *ieSi38*[*sun-1p*::*tir1*::*mRuby*::*sun-1* 3' UTR, *Cbr-unc-119* (+)] IV | This paper | CA1416 |
| *C. elegans*: *plk-2*(*ie108*[*plk-2*::*aid*::3xFLAG]), *zhp-2*(*ie107*[*zhp-2*::HA]) I; *mels8*[*pie-1p*::GFP::*cosa-1*, *unc-119*(+)] II; *ieSi38*[*sun-1p*::*tir1*::*mRuby*::*sun-1* 3'UTR, *Cbr-unc-119*(+)] IV | Zhang et al., 2018 | CA1429 |
| *C. elegans*: *mels8*[*pie-1p*::GFP::*cosa-1*, *unc-119*(+)] II; *ieSi38*[*sun-1p*::*tir1*::*mRuby*::*sun-1* 3' UTR, *Cbr-unc-119* (+)] IV; *chk-2*(*ie121*[HA::*aid*::*chk-2*]) V | This paper | CA1539 |
| *C. elegans*: *mels8*[*pie-1p*::GFP::*cosa-1*, *unc-119*(+)] II; *him-8*(*ie122*), *ieSi38*[*sun-1p*::*tir1*::*mRuby*::*sun-1* 3' UTR, *Cbr-unc-119* (+)] IV; *chk-2*(*ie121*[HA::*aid*::*chk-2*]) V | This paper | CA1540 |
| *C. elegans*: *plk-2*(*tm1395*) I; *mels8*[*pie-1p*::GFP::*cosa-1*, *unc-119*(+)] II; *ieSi38*[*sun-1p*::*tir1*::*mRuby*::*sun-1* 3' UTR, *Cbr-unc-119* (+)] IV; *chk-2*(*ie121*[HA::*aid*::*chk-2*]) V | This paper | CA1541 |
| *C. elegans*: *plk-2*(*ie50*[*plk-2*::*mRuby*] I??; *mels8*[*pie-1p*::GFP::*cosa-1*, *unc-119*(+)] II; *ieSi38*[*sun-1p*::*tir1*::*mRuby*::*sun-1* 3' UTR, *Cbr-unc-119* (+)] IV; *syp-1*(*ie123*[*syp-1* T452A]), *chk-2*(*ie121*[HA::*aid*::*chk-2*]) V | This paper | CA1542 |
| *C. elegans*: *plk-2*(*ie126*[*plk-2*::*aid*::HA]) I; *ieSi64*[*gld-1p*::*tir1*::*mRuby*::*gld-1* 3' UTR, *Cbr-unc-119* (+)] II; *cosa-1*(*ie124*[3xFLAG::*cosa-1*], *plk-1*(*ie125*[*plk-1*::*aid*::HA]) III | This paper | CA1543 |
| *C. elegans*: *plk-2*(*ie50*[*plk-2*::*mRuby*] I; *mels8*[*pie-1p*::GFP::*cosa-1*, *unc-119*(+)] II; *mpk-1*(*ie54*[*mpk-1*::*aid*::3xFLAG]) III; *ieSi38*[*sun-1p*::*tir1*::*mRuby*::*sun-1* 3' UTR, *Cbr-unc-119* (+)] IV; *syp-1*(*ie123*[*syp-1* T452A]) V | This paper | CA1544 |
| *C. elegans*: *ieSi64[gld-1p::tir1::mRuby::gld-1 3' UTR, Cbr-unc-119 (+)] II*; *cosa-1(ie124[3xFLAG::cosa-1] III* | This paper | CA1545 |
| *C. elegans*: *mels8[pie-1p::GFP::cosa-1, unc-119(+)] II*; *ieSi38[sun-1p::tir1::mRuby::sun-1 3' UTR, Cbr-unc-119 (+)] IV*; *chk-2(ie127[HA::chk-2]) V* | This paper | CA1546 |
| *C. elegans*: *plk-2(ie50[plk-2::mRuby] I??; mels8[pie-1p::GFP::cosa-1, unc-119(+)] II*; *ieSi38[sun-1p::tir1::mRuby::sun-1 3' UTR, Cbr-unc-119 (+)] IV??*; *syp-1(ie123[syp-1 T452A])*, *chk-2(ie127[HA::chk-2]) V* | This paper | CA1547 |
| *C. elegans*: *mels8[pie-1p::GFP::cosa-1, unc-119(+)] II*; *ieSi38[sun-1p::tir1::mRuby::sun-1 3' UTR, Cbr-unc-119 (+)] IV*; *chk-2(ie128[HA::chk-2 S116A]) V* | This paper | CA1548 |
| *C. elegans*: *mels8[pie-1p::GFP::cosa-1, unc-119(+)] II*; *ieSi38[sun-1p::tir1::mRuby::sun-1 3' UTR, Cbr-unc-119 (+)] IV*; *chk-2(ie129[HA::chk-2 S116A])/WT V* | This paper | CA1549 |
| *C. elegans*: *mels8[pie-1p::GFP::cosa-1, unc-119(+)] II*; *ieSi38[sun-1p::tir1::mRuby::sun-1 3' UTR, Cbr-unc-119 (+)] IV*; *chk-2(ie130[HA::chk-2 S116D])/WT V* | This paper | CA1550 |
| *C. elegans*: *mels8[pie-1p::GFP::cosa-1, unc-119(+)] II*; *ieSi38[sun-1p::tir1::mRuby::sun-1 3' UTR, Cbr-unc-119 (+)] IV*; *chk-2(ie131[HA::chk-2 T120A])/WT V* | This paper | CA1551 |
| *C. elegans*: *mels8[pie-1p::GFP::cosa-1, unc-119(+)] II*; *ieSi38[sun-1p::tir1::mRuby::sun-1 3' UTR, Cbr-unc-119 (+)] IV*; *chk-2(ie132[HA::chk-2 T120A])/WT V* | This paper | CA1552 |
| *C. elegans*: *mels8[pie-1p::GFP::cosa-1, unc-119(+)] II*; *ieSi38[sun-1p::tir1::mRuby::sun-1 3' UTR, Cbr-unc-119 (+)] IV*; *chk-2(ie133[HA::chk-2 T120D])/WT V* | This paper | CA1553 |
| *C. elegans*: *mels8[pie-1p::GFP::cosa-1, unc-119(+)] II*; *ieSi38[sun-1p::tir1::mRuby::sun-1 3' UTR, Cbr-unc-119 (+)] IV*; *chk-2(ie134[HA::chk-2 T120D])/WT V* | This paper | CA1554 |
| *C. elegans*: *mels8[pie-1p::GFP::cosa-1, unc-119(+)] II*; *ieSi38[sun-1p::tir1::mRuby::sun-1 3' UTR, Cbr-unc-119 (+)]*, *plk-3(ie135)*, *him-8(ie122) IV*; *chk-2(ie121[HA::aid::chk-2]) V* | This paper | CA1555 |
| *C. elegans*: *plk-2(ie126[plk-2::aid::HA]) I*; *ieSi64[gld-1p::tir1::mRuby::gld-1 3' UTR, Cbr-unc-119 (+)] II*; *cosa-1(ie124[3xFLAG::cosa-1]*, *plk-1(ie125[plk-1::aid::HA]) III*; *plk-3(ie136) IV* | This paper | CA1556 |
| *C. elegans*: *mels8[pie-1p::GFP::cosa-1, unc-119(+)] II*; *ieSi38[sun-1p::tir1::mRuby::sun-1 3' UTR, Cbr-unc-119 (+)] IV*; *him-5(ok1896)*, *chk-2(ie121 [HA::aid::chk-2]) V* | This paper | CA1557 |
| *C. elegans*: *mels8[pie-1p::GFP::cosa-1, unc-119(+)] II*; *ieSi38[sun-1p::tir1::mRuby::sun-1 3' UTR, Cbr-unc-119 (+)] IV*; *pas-1(ie185[pas-1::aid::3xFLAG]) V* | This paper | CA1558 |
| *C. elegans*: *mels8*[*pie-1p::GFP::cosa-1, unc-119(+)*] II; *ieSi38*[*sun-1p::tir1::mRuby::sun-1 3' UTR, Cbr-unc-119 (+)*] IV; *pas-1*(*ie185*[*pas-1::aid::3xFLAG*]), *chk-2*(*ie186* [ALFA::*chk-2*]) V | This paper | CA1559 |
| *C. elegans*: *plk-2*(*ie126*[*plk-2::aid::HA*]) I; *ieSi64*[*gld-1p::tir1::mRuby::gld-1 3' UTR, Cbr-unc-119 (+)*] II; *cosa-1*(*ie124*[*3xFLAG::cosa-1*], *plk-1*(*ie125*[*plk-1::aid::HA*]) III; *chk-2*(*ie187* [ALFA::*chk-2*]) V | This paper | CA1628 |
| *C. elegans*: *plk-2*(*ie126*[*plk-2::aid::HA*]) I; *ieSi64*[*gld-1p::tir1::mRuby::gld-1 3' UTR, Cbr-unc-119 (+)*] II; *cosa-1*(*ie124*[*3xFLAG::cosa-1*], *plk-1*(*ie125*[*plk-1::aid::HA*]) III; *pas-1*(*ie188*[*pas-1::aid::3xFLAG*]), *chk-2*(*ie187* [ALFA::*chk-2*]) V | This paper | CA1629 |

